# Fasting is required for many of the benefits of calorie restriction in the 3xTg mouse model of Alzheimer’s disease

**DOI:** 10.1101/2024.09.19.613904

**Authors:** Reji Babygirija, Jessica H. Han, Michelle M. Sonsalla, Ryan Matoska, Mariah F. Calubag, Cara L. Green, Anna Tobon, Chung-Yang Yeh, Diana Vertein, Sophia Schlorf, Julia Illiano, Yang Liu, Isaac Grunow, Michael J. Rigby, Luigi Puglielli, David A. Harris, John M. Denu, Dudley W. Lamming

## Abstract

Caloric restriction (CR) is a widely recognized geroprotective intervention that slows or prevents Alzheimer’s disease (AD) in animal models. CR is typically implemented via feeding mice a single meal per day; as CR mice rapidly consume their food, they are subject to a prolonged fast between meals. While CR has been shown to improve metabolic and cognitive functions and suppress pathological markers in AD mouse models, the specific contributions of fasting versus calorie reduction remains unclear. Here, we investigated the contribution of fasting and energy restriction to the beneficial effects of CR on AD progression. To test this, we placed 6-month-old 3xTg mice on one of several diet regimens, allowing us to dissect the effects of calories and fasting on metabolism, AD pathology, and cognition. We find that energy restriction alone, without fasting, was sufficient to improve glucose tolerance and reduce adiposity in both sexes, and to reduce Aβ plaques and improve aspects of cognitive performance in females. However, we find that a prolonged fast between meals is necessary for many of the benefits of CR, including improved insulin sensitivity, reduced phosphorylation of tau, decreased neuroinflammation, inhibition of mTORC1 signaling, and activation of autophagy, as well as for the full cognitive benefits of CR. Finally, we find that fasting is essential for the benefits of CR on survival in male 3xTg mice. Overall, our results demonstrate that fasting is required for the full benefits of a CR diet on the development and progression of AD in 3xTg mice, and suggest that both when and how much we eat influences the development and progress of AD.

## Introduction

The global population is rapidly graying; with this silver tsunami, the prevalence of Alzheimer’s disease (AD) is increasing and by 2050, 13.8 million Americans are projected to be struggling with AD ^1,2^. Decades of research to discover treatments for AD have been largely unsuccessful as well as extremely expensive; identifying new and cost-effective interventions that can delay or prevent AD is therefore of critical importance.

Caloric restriction (CR), a reduction in caloric intake without malnutrition or starvation, is the most robust, non-pharmacological and reproducible intervention for extending lifespan in model organisms ^3^. CR slows the development of AD in multiple mouse models ^4–7^, as well as in squirrel monkeys ^8^. While the impact of CR on AD in humans is mostly unknown, short-term CR can improve memory ^9^. As few humans are prepared to engage in long-term CR, there is substantial interest in understanding the mechanisms underlying CR to harness its benefits without the need to subject individuals to an abstemious dietary regimen.

Many studies have investigated potential molecular mechanisms which may be engaged by a CR diet. To name just a few, CR decreases mammalian target of rapamycin complex 1 (mTORC1) signaling and activates autophagy. We recently asked a different question based on the realization that CR-fed mice have an unusual feeding pattern. While *ad libitum* (AL)-fed mice eat in multiple small bouts throughout the day, with most of the intake during the night, CR regimens are performed by feeding mice only once-per-day. As a result, CR mice binge eat their food within 1-2 hours^10,11^ and are subjected to a self-imposed ∼22 hour fast between meals. We recently developed a series of feeding regimens that enabled us to dissect the relative contribution of fasting from energy restriction in the CR diet paradigm. We found that in wild-type mice, the prolonged fast between meals was necessary for the beneficial effects of a CR diet on metabolic health, frailty, longevity, and cognition ^12^.

In the present study, we expanded upon this work and set out to define the relative importance of fasting to the benefits of CR in AD. We used the 3xTg mouse model of AD, which expresses familial human isoforms of APP (APPSwe), Tau(tauP301L), and Presenilin (PS1M146V), and exhibits both Aβ and tau pathology as well as cognitive deficits ^13^. 3xTg mice have been widely used to study geroprotective interventions, and CR has clearly beneficial effects on AD pathology and cognition in this model ^4^. We placed 3xTg mice on a series of dietary regimens with similar compositions and investigated whether the benefits of CR on AD are mediated by energy restriction, fasting, or a combination of both.

Here, we find that fasting is not only required for many of the metabolic benefits of CR in the 3xTg mouse, but that it is also required for many of the molecular effects of CR, for CR-induced improvements in tau pathology and neuroinflammation, and for the full cognitive benefits of CR. Surprisingly, fasting is not required for CR-induced reductions in brain Aβ plaques accumulation in females, while CR had no effect on Aβ plaques in males. These results suggest that many of the beneficial effects of CR on memory, especially in males, may result from reduced tau pathology or altered molecular signaling, and not from the reduction in plaques. Our results demonstrate that fasting is a critical component of the benefits of CR on AD suggesting that fasting alone or fasting-mimicking diets might help to prevent or delay AD in humans without requiring reductions in energy intake.

## Results

### CR has sex-specific effects on body composition and energy balance

We randomized 6-month-old male and female 3xTg mice to one of three groups of equivalent body weight then placed each on one of three dietary regimens: *Ad* libitum (AL) - mice with free access to a normal rodent diet (Envigo Global 2018; **Supplementary Table 1**), Diluted *ad libitum* (DL) - mice with free access to Envigo Global 2018 diluted 50% with indigestible cellulose (TD.170950; **Supplementary Table 1**); equivalent to 30% restriction of calories without imposing fasting, and caloric restriction (CR) - mice fed a normal rodent diet (Envigo Global 2018) once per day in the morning, with 30% restriction of calories relative to AL-fed mice and with prolonged inter-meal fasting.

We followed the mice longitudinally for 9 months, tracking their body weight and determining their body composition at the beginning and the end of the experiment (**Fig. 1A**). As we expected, neither CR nor DL-fed female 3xTg mice gained weight during the course of the experiment, while AL-fed females mice continued to gain weight (**Fig. 1B**); this was primarily the result of an effect of CR and DL-feeding on fat mass, which resulted in an overall reduction in adiposity (**Figs. 1C-E**). We tracked food consumption rigorously and observed that both CR- and DL-fed mice had comparable calorie intake (**Fig. 1F**).

**Figure 1:**
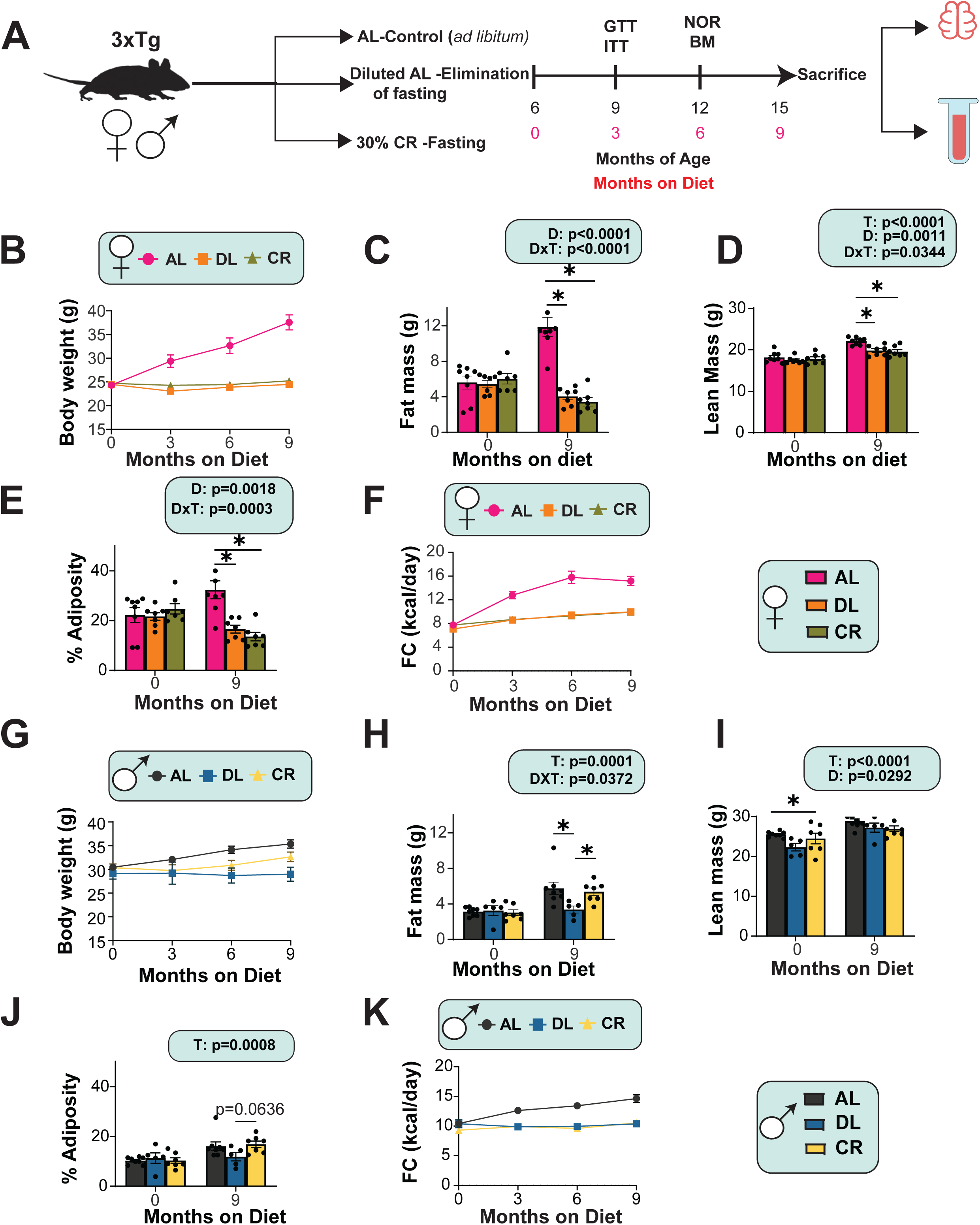
Metabolic health outcomes of 3xTg-AD mice on different feeding regimens. (A) Experimental design: Six-month-old female and male 3xTg-AD mice were placed on either ad libitum (AL), Diluted AL or Caloric Restricted (CR) diet and phenotyped over the course of the next 9 months. (B-E) The body weight (B) of female mice was followed over the course of the experiment, fat mass (C) and lean mass (D) was determined at the start and end of the experiment, and the adiposity (E) was calculated. (B-E) n=8 AL, n=7 DL and n=7 CR fed 3xTg biologically independent mice. (F) Food consumption of female mice at 0,3,6 and 9 months of age; n=8 AL, n=7 DL and n=7 CR fed 3xTg biologically independent mice. (G-J) The body weight (G) of male mice was followed over the course of the experiment, fat mass (H) and lean mass (I) was determined at the start and end of the experiment, and the adiposity (J) was calculated. (G-J) n=8 AL, n=5 DL, n=7 CR 3xTg biologically independent mice. (K) Food consumption of male mice at 0, 3, 6 and 9 months of age; n=8 AL, n=5 DL, n=7 CR 3xTg biologically independent mice. (C-E, H-J) Statistics for the overall effect of diet and time represent the p value from a two-way analysis of variance (ANOVA); *p<0.05, Sidak’s post-test examining the effect of parameters identified as significant in the two-way ANOVA. Data represented as mean ± SEM.

3xTg males had an initial response in body weight to the diet regimens that was similar to the 3xTg females, but as the experiment continued, the CR-fed mice started to weigh more than the DL-fed males (**Fig. 1G**), and by the end of the experiment AL- and CR-fed mice had comparable levels of fat mass, lean mass, and adiposity, while DL-fed males had reduced fat mass and adiposity relative to both AL and CR-fed males (**Figs. 1H-J**). As with females, CR and DL-fed 3xTg males had comparable reductions in calorie intake relative to AL-fed males (**Fig. 1K**).

CR-fed mice engage in rapid lipogenesis following refeeding, and then sustain themselves with these stored lipids ^14^. We can see these shifts in fuel utilization by using metabolic chambers to determine substrate utilization by analyzing the respiratory exchange ratio (RER), which is derived from oxygen consumption and carbon dioxide production. A high RER value signifies the utilization of carbohydrates for energy production or lipogenesis, while a value nearing 0.7 indicates that lipids serve as the primary energy source. As we anticipated, both male and female CR-fed 3xTg mice have a distinct RER curve from AL and DL-fed mice, reflecting the rapid induction of lipogenesis following feeding and a subsequent switch to utilizing lipids as the primary energy source (**Figs. 2A-B, D-E**). CR-fed 3xTg mice had decreased energy expenditure relative to AL-fed mice in both sexes; however, while DL-fed males had lower energy expenditure, this effect did not reach statistical significance (**Figs. 2C and 2F**).

**Figure 2:**
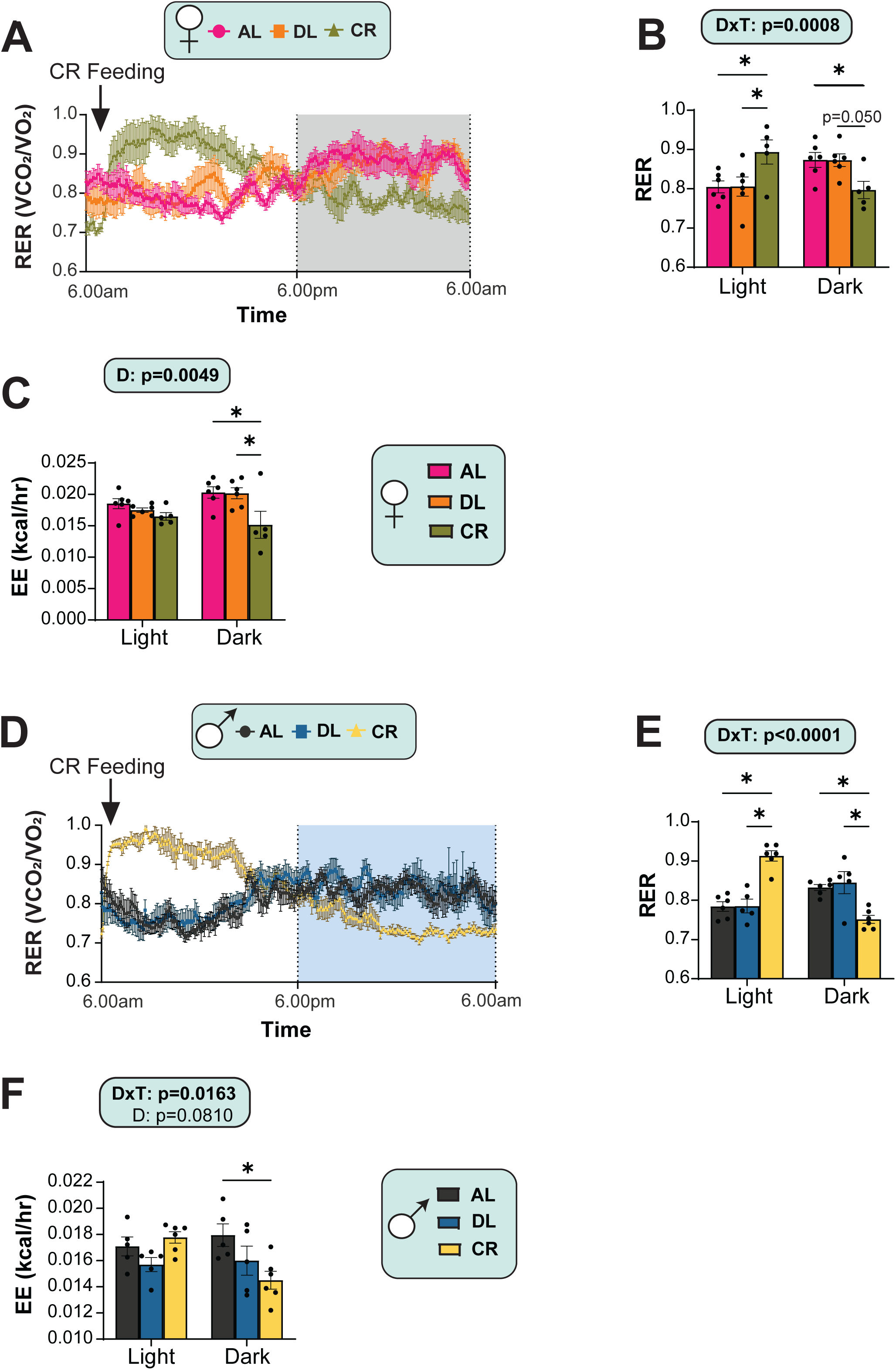
Distinct fuel utilization patterns of 3xTg-AD mice under reduced caloric intake conditions. (A-F) Metabolic chambers were used to determine fuel source utilization and energy expenditure 24 hours in female (A-C) and male (D-F). Six-month-old female and male 3xTg-AD mice were placed on either ad libitum (AL), Diluted AL or Caloric Restricted (CR) diet and phenotyped over the course of the next 9 months. (A) Respiratory exchange ratio (RER) over 24hrs in females (B) RER during light and dark cycles in females. (C) Energy expenditure (EE) normalized to body weight in females (D) RER over 24 hrs in males (E) RER during light and dark cycles in males (F) EE normalized to body weight in males. (A-C) n=6 AL, n=6 DL and n=5 CR fed biologically independent 3xTg female mice were used. (D-F) n=5 AL, n=5 DL and n=6 CR fed biologically independent 3xTg male mice were used. (B-C, E-F) Statistics for the overall effect of diet and time represent the p value from a two-way ANOVA; *p<0.05, Sidak’s post-test examining the effect of parameters identified as significant in the two-way ANOVA. Data represented as mean ± SEM.

Improved glucose tolerance and insulin sensitivity is a hallmark of CR in mammals. We performed glucose and insulin tolerance tests (GTT and ITT, respectively), timing the assays such that mice in all groups were fasted (22hrs) for similar lengths of time. As anticipated, both CR and DL-fed female 3xTg mice had improved glucose tolerance compared to the AL fed group (**Fig. 3A**). Strikingly, in agreement with our previous results in C57BL/6J mice, insulin sensitivity as assessed by an intraperitoneal ITT was significantly improved only in CR-fed mice, and not in the DL-fed group (**Fig. 3B**). We observed similar, if stronger, effects of CR and DL on GTT and ITT in male mice (**Figs. 3C-D**).

**Figure 3:**
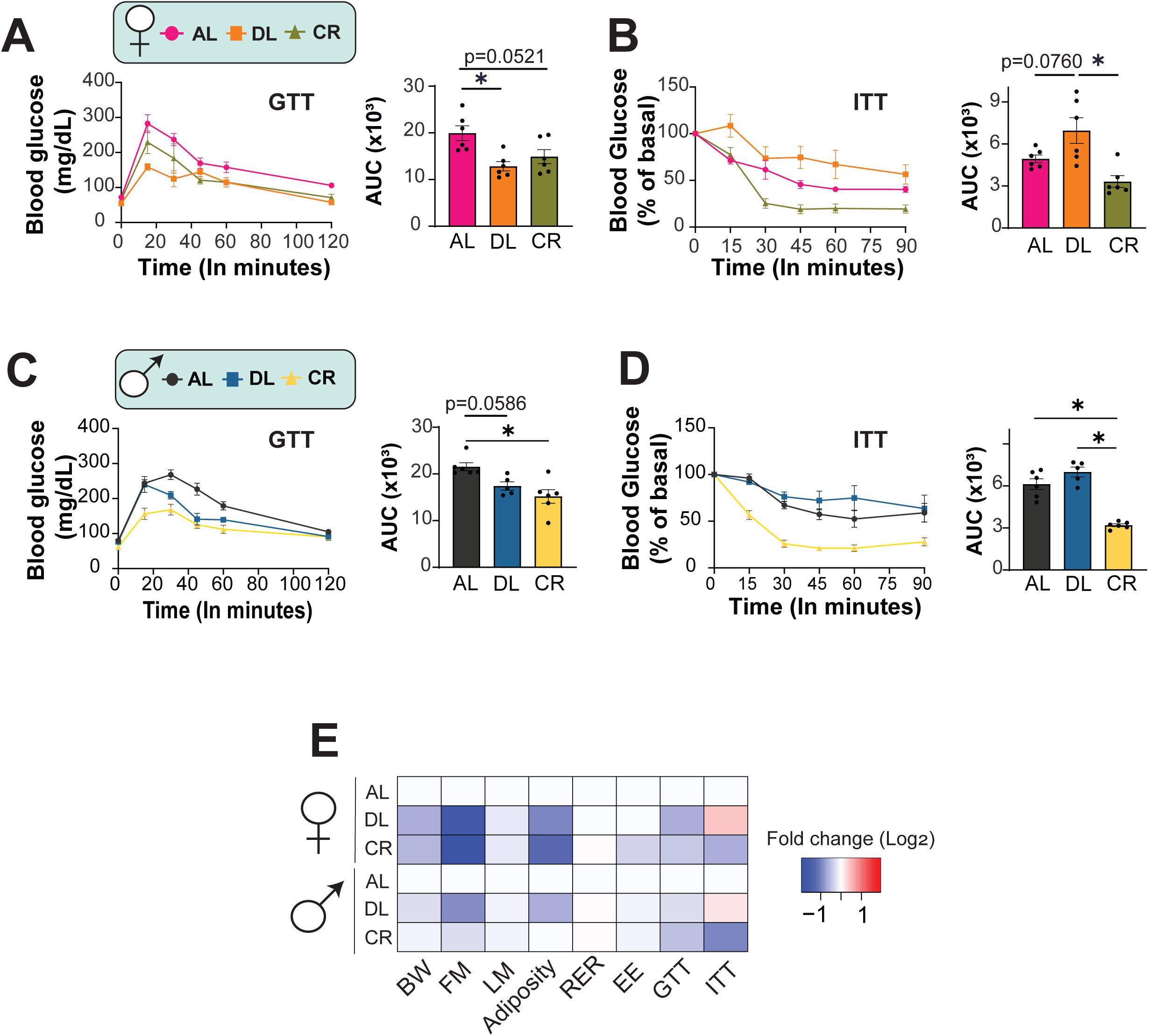
Fasting is required for CR-induced insulin sensitivity in both female and male 3xTg-AD mice. (A-B) Glucose (A) and insulin (B) tolerance tests were performed in female mice after three months on AL, DL or CR fed diets. (A) GTT: n=6 3xTg biologically independent mice per group. (B) ITT: n=6 3xTg biologically independent mice per group. (C-D) Glucose (C) and insulin (D) tolerance tests were performed in male mice after three months on AL, DL or CR fed diets. (A) GTT: n=6 AL, n=5 DL and n=6 CR fed 3xTg biologically independent mice (B) ITT: n=6 AL, n=5 DL and n=6 CR fed biologically independent mice. (E) Heat map representation of all the metabolic parameters in 3xTg female and male mice. Color represents the log_2_ fold-change vs. AL fed mice. (A-D) *p<0.05, Tukey post-test examining the effect of parameters identified as significant in the one-way ANOVA. Data represented as mean ± SEM. AUC, Area Under the Curve.

Overall, we found that CR and DL-fed 3xTg mice of both sexes showed similar improvements in metabolic parameters including weight, body composition and energy balance. However, improvements in insulin sensitivity were observed only in CR-fed mice. Thus, as observed in wild-type mice, CR-induced improvements in insulin sensitivity require fasting in 3xTg mice of both sexes (**Fig. 3E**).

### Sex-specific effects of energy restriction with and without fasting on metabolism

To investigate potential differences and overlaps on the effects of energy restriction with and without fasting, we performed a targeted metabolomics analysis of plasma and brain of AL, CR, and DL-fed mice of both sexes. Our analysis targeted ∼50 metabolites across three molecular groups based on their chemical structures: amino acids, carbohydrates, and nucleosides/nucleotides.

Principal Component Analysis (PCA) revealed distinct metabolic signatures in the plasma of AL, CR, and DL-fed 3xTg females (**Fig. 4A**). Plotting the 47 targeted plasma metabolites illustrates the shared and distinct metabolic signatures induced by CR and DL diets relative to AL-fed females (**Fig. 4B, Table S2-S3**). Among the significantly altered metabolites, we observed significant CR-induced increases in guanosine, leucine, and isoleucine, as well as CR-induced reductions in uridine. These changes were specific to CR and were not induced by a DL diet; the only significantly altered metabolite that was shared in both CR and DL-fed mice was glucose-6-phosphate, which was decreased in the plasma of both groups (**Figs. S1A-C**). Intriguingly, these patterns are sex-specific and were not observed among 3xTg males (**Figs. S2A-B, Table S4-S5**).

**Figure 4:**
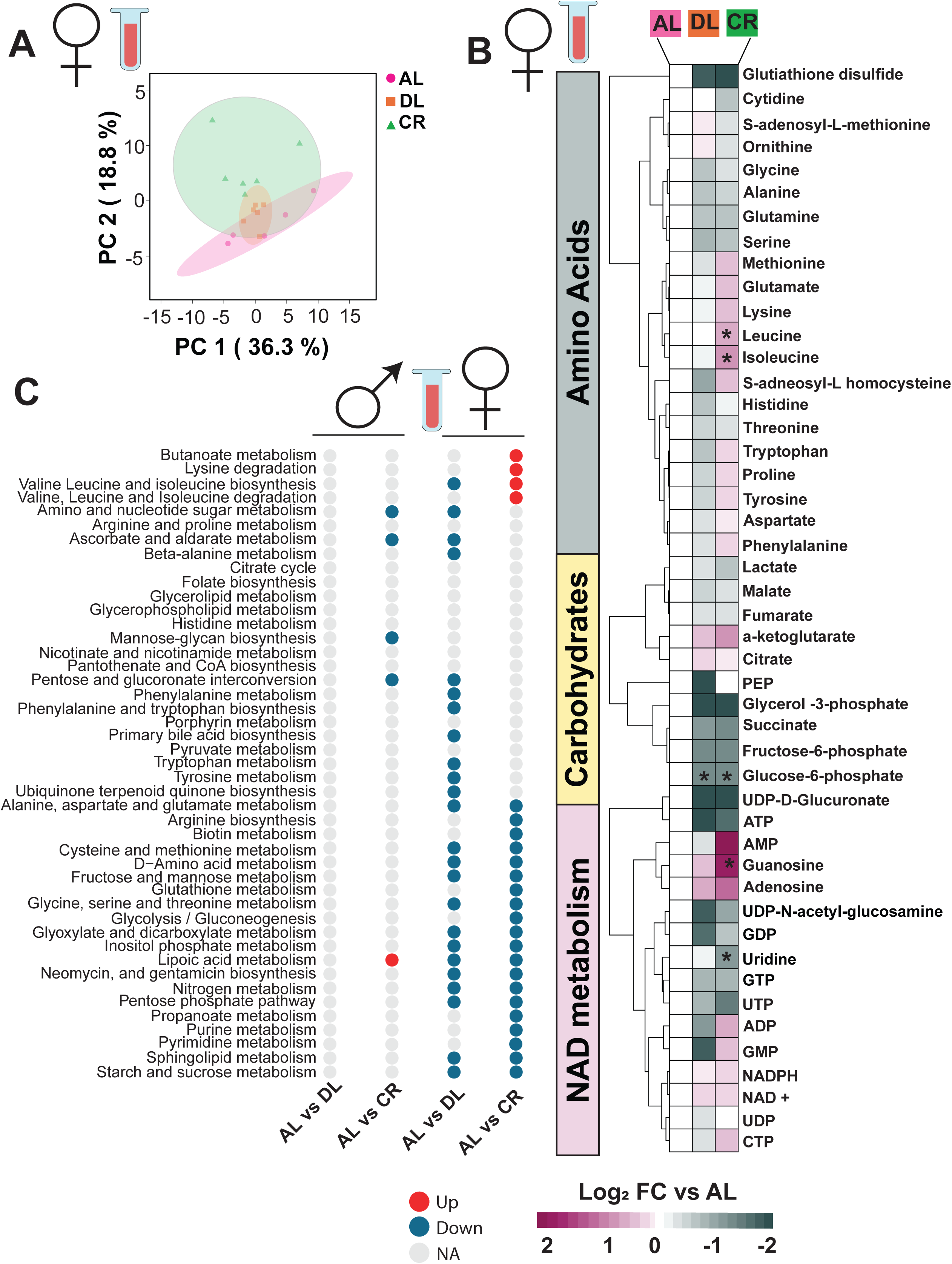
Plasma metabolome displays sex specific effects of fasting and energy restriction in 3xTg-AD mice. Targeted metabolomics were performed on the plasma of female 3xTg mice fed AL, Diluted AL and CR diets (n = 6) biologically independent mice per diet. (A) Principal Component Analysis (PCA) of plasma metabolites from 3xTg females fed on an AL, DL or CR diets (B) Heatmap of 47 targeted metabolites, represented as log2-fold change vs. AL-fed mice. * Symbol represents a significant difference versus AL mice (FDR, q < 0.05). (C) Significantly up and down regulated pathways for each sex and diet were determined using metabolite set enrichment analysis (MSEA).

Shifts in the levels of multiple metabolites within a single pathway can provide more information about regulatory processes than changes within levels of a single metabolite. We utilized Metabolite Set Enrichment Analysis (MSEA) to identify the distinct and shared pathways altered by each diet regimen. Our analysis identified multiple metabolic pathways that were altered in both 3xTg CR-fed and DL-fed females, including downregulation of pathways involving the metabolism of amino acids including “Cysteine and methionine metabolism”, “Glycine, serine and threonine metabolism,” and “Alanine, aspartate and glutamate metabolism” (**Fig. 4C, Table S6**). There were also pathways altered specifically by each diet in 3xTg females; CR-fed females showed upregulation of pathways including “Butanoate metabolism”, “Lysine degradation,” and “Valine, leucine, and isoleucine degradation.” Interestingly, DL-fed females had no significantly upregulated pathways, and indeed CR-fed females had upregulation of “Valine, leucine, and isoleucine biosynthesis,” while DL-fed females had downregulation of this same pathway. In contrast, very few pathways were significantly altered in CR-fed 3xTg males, and no pathways were significantly altered in DL-fed 3xTg males (**Fig. 4C, Table S6**).

We also examined the metabolic signature in the whole brain of 3xTg male and female mice on each diet regimen. PCA on the identified 58 brain metabolites showed extensive overlap between female 3xTg mice fed any of the three diets (**Fig. 5A, Table S7**). In particular, we were surprised to find that the effect of DL and CR diets on brain metabolites was virtually identical; a total of 21 metabolites were significantly altered by either the CR or DL diets or were altered in both. This included elevations in the levels of many essential and non-essential amino acids, including methionine and S-Adenosyl homocysteine (SAH), a metabolite in the cysteine and methionine metabolism pathway (**Figs. 5B, S3A-C and Table S8**). Reflecting this high degree of overlap, MSEA found many biological pathways that were significantly affected by both DL and CR feeding in the brains of 3xTg female mice; pathways uniquely upregulated in CR-fed females included “Propanoate metabolism,” “Folate biosynthesis,” “Pentose phosphate pathway,” “Lipoic acid metabolism,” and “Glycerolipid metabolism” (**Fig. 5C, Table S9**).

**Figure 5:**
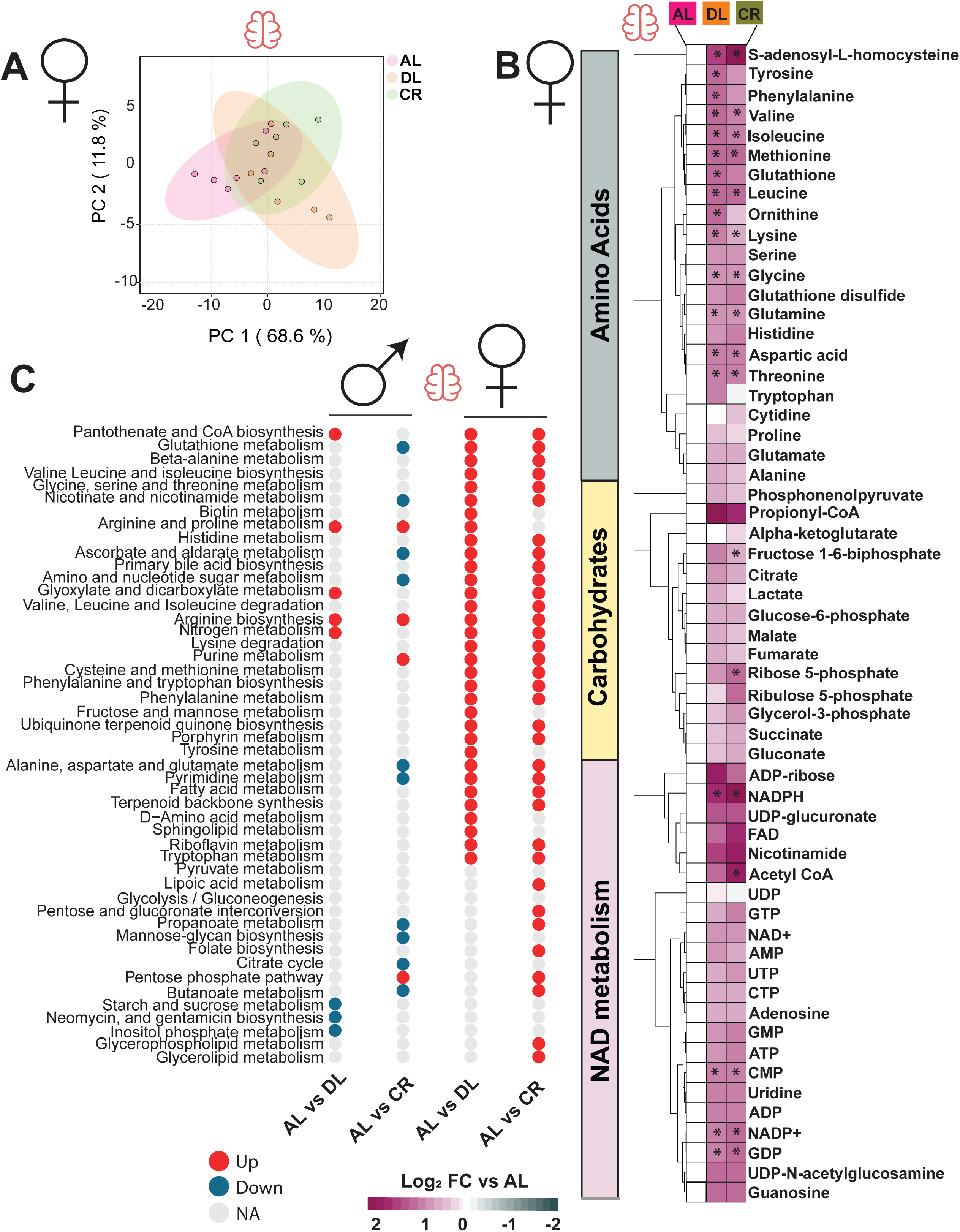
Overlapping brain metabolome profile in female 3xTg-AD mice under energy restriction and once-per-day CR regimen. Targeted metabolomics were performed on the brain of female 3xTg mice fed AL, Diluted AL and CR diets (n = 6) biologically independent mice per diet. (A) Principal Component Analysis (PCA) of brain metabolites from 3xTg females fed on an AL, DL or CR diets (B) Heatmap of 59 targeted metabolites, represented as log2-fold change vs. AL-fed mice. * Symbol represents a significant difference versus AL mice (FDR, q < 0.05) based on Tukey’s test post one-way ANOVA. (C) Significantly up and down regulated pathways for each sex and diet were determined using metabolite set enrichment analysis (MSEA).

As in the brain, we observed many fewer changes in 3xTg males fed either the CR or DL diets than in females; however, PCA showed that CR was clearly distinct from both the AL and DL-fed groups (**Fig. S4A and Table S10**). While there were similar trends in the effect of CR and DL diets on NAD metabolism (**Fig. S4B, Table S11**), only NADP+ and ribulose-5 phosphate were significantly altered, and only in CR-fed 3xTg males (**Fig. S4C-D**). Among the significantly altered pathways in both DL and CR-fed 3xTg males were” Arginine and proline metabolism” and “Arginine biosynthesis” which may reflect the progression of neurodegeneration ^15,16^ (**Fig. 5C**). Notably, “Arginine biosynthesis” was similarly upregulated in both CR and DL-fed female 3xTg mice, the only pathway so similarly regulated in all four groups. Notably, the effect of CR and DL feeding on “Cysteine and Methionine metabolism” was observed in both plasma and brain samples of female, but not male, 3xTg mice, with similar effects on most of the metabolites detected despite the pathway being upregulated in the brain and downregulated in the plasma (**Figs. S5A-B**).

### Fasting is required for CR-induced improvements in tau hyperphosphorylation and neuroinflammation, but not for CR-induced improvements in Aβ plaque deposition

To investigate the role of fasting in the ability of CR to attenuate AD pathology, we evaluated several pathological hallmarks of AD, including phosphorylation of tau, amyloid beta (Aβ) plaque deposition, and gliosis. In earlier studies ^17^, we did not see any visible plaques in 12-month-old 3xTg mice, and we therefore evaluated AD pathology in in the cortex and hippocampus of both male and female 3xTg mice at 15 months of age. Brain sections from 15-month-old 3xTg mice females were immune stained and visualized using 3,3’-diaminobenzidine (DAB) (**Fig. 6A**). We observed significant plaque deposition in AL-fed females which was significantly diminished by both DL and CR, as shown by the reduced plaque area in both the cortex and hippocampus of female 3xTg mice fed these diets (**Fig. 6A**). However, upon immunoblotting for phosphorylated tau at phosphorylation site Thr231, we observed that only CR-fed mice had reduced tau phosphorylation, which reached statistical significance as compared to DL-fed mice (**Fig. 6B**). Furthermore, fluorescent immunostaining combined with quantitative analysis of cortical p-Tau Thr231 in female 3xTg mice also showed significant decrease in p-Tau only in CR-fed group **(Fig. 6C).**

**Figure 6:**
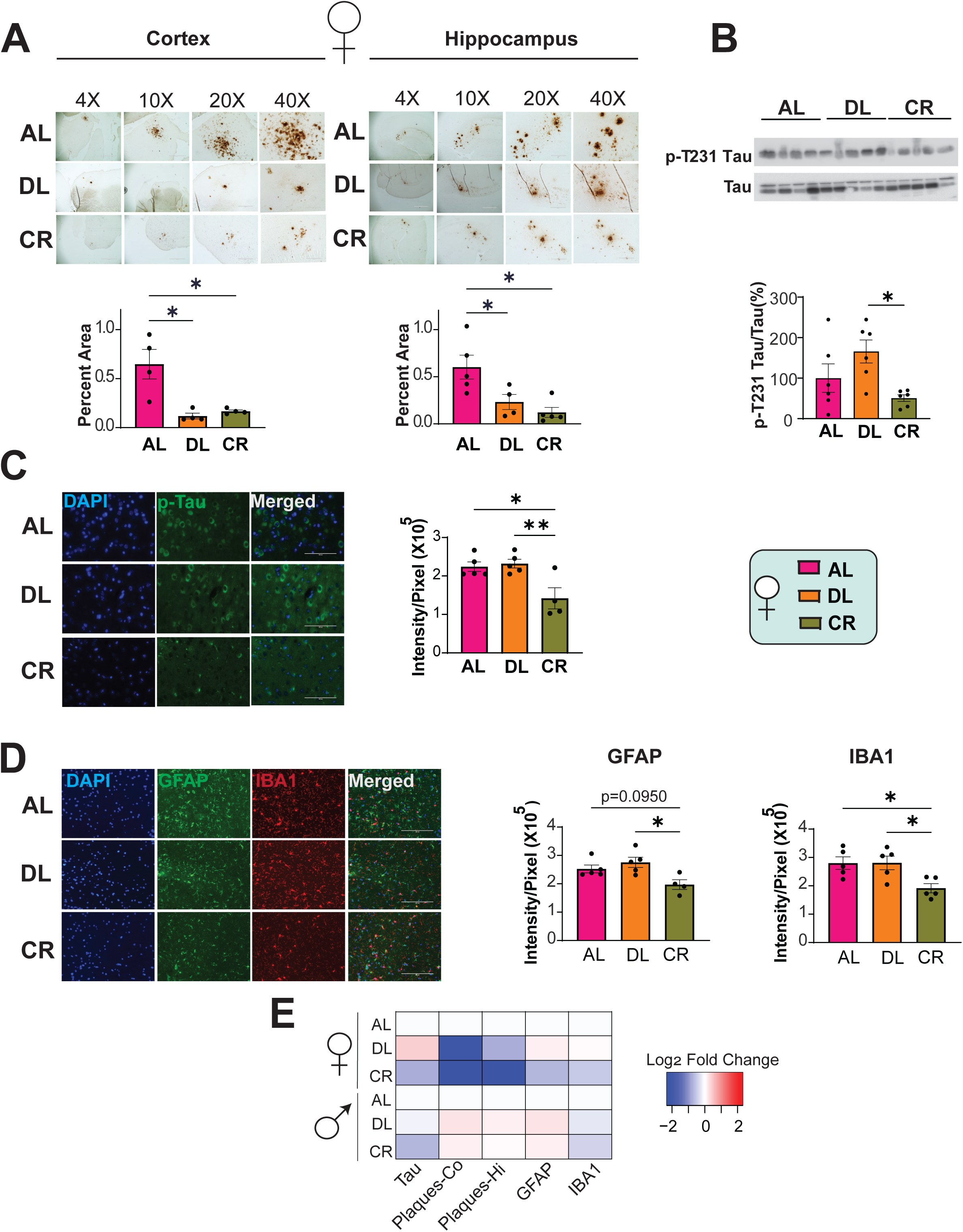
Fasting required for mitigating hyperphosphorylation of tau, but not Aβ plaques in female 3xTg-AD mice. (A-C) Analysis of AD neuropathology in female 3xTg mice fed on AL, Diluted AL or CR diet from 6-15 months of age. (A) Representative plaque images of DAB staining with 6E10 antibody in the cortex and hippocampus of female 3xTg mice. 4x, 10x, 20x and 40x magnification shown; scale bar in the 4x image is 1000 µM, 10x image is 400µM, 20x is 200 µM and 40x is 100 µM. Quantification of plaque area in females shown under each representative blots from cortex and hippocampus respectively. For cortex n=4 biologically independent mice per group and for hippocampus n=5 AL, n=4 DL and n=5 CR fed 3xTg biologically independent mice. (B) Western blot analysis of phosphorylated Th231 Tau in female 3xTg mice (n=6 biologically independent mice/group). (C) Representative immunofluorescence images in the cortex of 3xTg females stained with p-Tau Thr231 antibody (AT180), 40x magnification shown; scale bar 100 µM. Quantitative analysis of fluorescence intensity. n=5 AL, n=5 DL, n=4 CR fed 3xTg biologically independent mice. (D) Immunostaining and quantification of 5 μm paraffin-embedded brain slices for astrocytes (GFAP) and microglia (Iba1) in female 3xTg mice. 20x magnification shown; Scale bar is 200 µM. n=5 biologically independent mice/group. (A-C) *p<0.05, Tukey post-test examining the effect of parameters identified as significant in the one-way ANOVA. (E) Heat map representation of the neuropathological findings in female and male 3xTg mice; log_2_ fold-change relative to AL-fed mice of each sex. Data represented as mean ± SEM.

As expected, the plaque-burdened AL-fed 3xTg females had significant neuroinflammation with higher levels of activated astrocytes (GFAP-positive) and microglia (IBA1-positive). CR-fed 3xTg females showed a significant reduction in both GFAP- and IBA1-positive cells (**Fig. 6D**). In contrast, despite a significant reduction in plaque burden, DL-fed 3xTg females did not show a reduction in the number of either GFAP or IBA1-positive cells (**Fig. 6D**).

Interestingly, in males, we did not observe significant plaque deposition as observed in females (**Fig. S6A**). However, CR-fed 3xTg males did showed a significant reduction in phosphorylated tau (**Fig. S6B and S6C**), as well as a significant reduction in IBA1 positive cells as compared to the AL-fed mice (**Fig. S6C**). Taken together, these results suggest that fasting is essential for reducing hyperphosphorylation of tau and gliosis in both females and males, independent of plaque deposition (**Fig. 6E**).

### CR and DL have distinct effects on mTORC1 signaling and autophagy

mTORC1 signaling has been extensively studied in the brains from AD mouse models, revealing mTORC1 activation in these brains ^18,19^. It is widely believed that CR decreases mTORC1 signaling, which would result in the activation of autophagy while reducing overall biosynthesis, decreasing protein translation as well as lipid synthesis ^20^. We therefore investigated how once-per-day CR and energy restriction alone impact mTORC1 signaling in the brains of the AD mice.

We performed immunoblotting in brain lysates to assess the phosphorylation of mTORC1 substrates p-S240/S244 S6 and T37/S46 4E-BP1. Compared to AL and DL-fed 3xTg females, we observed significantly decreased phosphorylation of both substrates in CR-fed 3xTg females (**Figs. 7A-C**).

**Figure 7:**
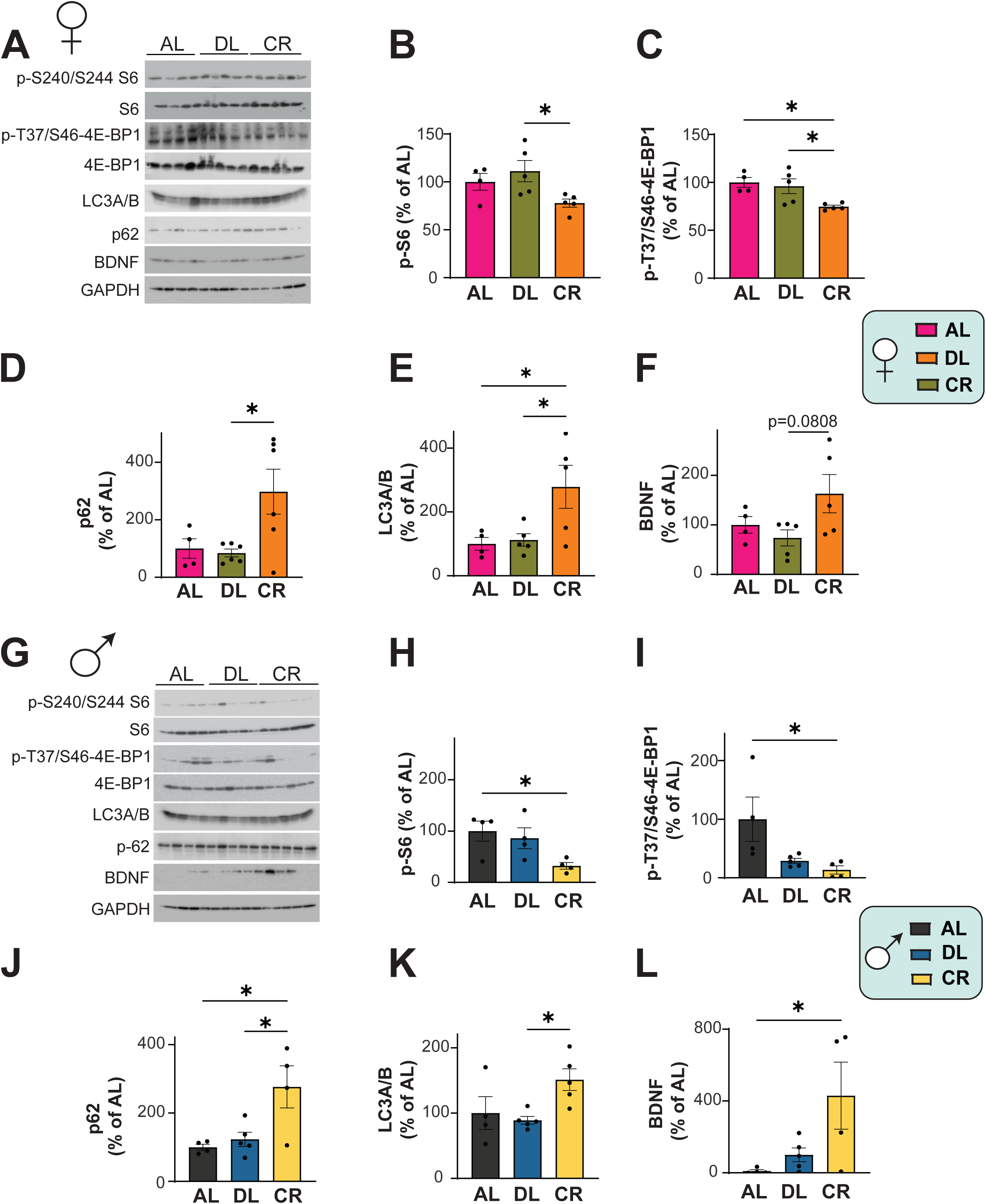
Differential effects of feeding regimens on brain mTORC1 signaling and autophagy activation. (A-L) The phosphorylation of S6 and 4E-BP1, and the expression of p62, LC3A/B and BDNF was assessed by western blotting of whole brain lysates. (B, C) Quantification of the phosphorylation of p-S240/S244 S6 (B) and T37/S46 4E-BP1 (C), relative to expression of S6 and 4E-BP1, respectively in females. (D-F) Quantification of p62 expression (D) LC3A/B expression (E) and BDNF expression (F) relative to expression of GAPDH in females. (H, I) Quantification of the phosphorylation of p-S240/S244 S6 (H) and T37/S46 4E-BP1 (I), relative to expression of S6 and 4E-BP1, respectively in males. (J-L) Quantification of p62 expression (J) LC3A/B expression (K) and BDNF expression (L) relative to expression of GAPDH in males (B, C, E, F) n=4 AL, n=5 DL and n=5 CR fed 3xTg biologically independent female mice. (D) n=4 AL, n=6 DL and n=6 CR fed 3xTg biologically independent female mice (H-L) n=4 AL, n=5DL and n=5 CR fed 3xTg biologically independent male mice. (B-F, H-L) *p<0.05, Tukey post-test examining the effect of parameters identified as significant in the one-way ANOVA. Data represented as mean ± SEM. BDNF; Brain derived neurotrophic factor

We assessed autophagy by examining expression of the autophagy receptor p62 (sequestosome 1, SQSTM1) and the autophagosome marker LC3A/B. We found that CR-fed female mice exhibited increased expression of both p62, and LC3A/B compared to AL and DL-fed groups (**Figs. 7D-E**). These results indicate that fasting is critical for both CR-mediated suppression of mTORC1 signaling and CR-dependent induction of autophagy.

We also investigated the expression of brain-derived neurotrophic factor (BDNF), which is known for its neuroprotective effects; reduced BDNF levels are reported in the pathogenesis of AD ^21–29^. Moreover, BDNF has been shown to modulate mTORC1 signaling specifically in the neurons to promote synaptic plasticity and neuronal growth ^30,31^. We found that BDNF levels were increased in the CR-fed 3xTg females compared to DL-fed mice (p=0.0808), which could suggest that upregulation of BDNF confers neuroprotective effects in the once-a day fed CR group (**Fig. 7F**).

In males, we observed very similar results to those we found in females, with significant downregulation of p-S240/S244 S6 and T37/S46 4E-BP1 in the CR-fed group compared to the AL-fed group (**Figs. 7G-I**). While the DL-fed 3xTg males exhibited reduced phosphorylation of mTORC1 substrates compared to AL-fed males, these reductions were not statistically significant (**Figs. 7G-I**). We also observed significant activation of autophagy markers in the CR-fed groups compared to AL and DL-fed mice **(Fig. 7J-K).** Finally, BDNF expression was significantly upregulated in CR-fed 3xTg males, which aligns with the improvements in AD pathology, particularly the tau-specific improvements in the males (**Fig. 7L**).

### Fasting is required for rescuing hippocampal-dependent long-term memory in both sexes

We examined the contribution of fasting on CR-induced improvements in cognition by performing behavioral assays on 12-month-old 3xTg mice that had been fed either an AL, DL, or CR diet prior to sacrificing them for the histological assays described above. We examined performance of both sexes in a Barnes Maze (BM) and tested Novel Object Recognition (NOR).

In the BM assay, mice were required to locate an escape box placed at the target hole using spatial cues during days 1-4 of the acquisition phase. On days 5 and 12 of the BM assay short-term memory (STM) and long-term memory (LTM), respectively, were tested, evaluating how well the mice remembered the location of the target hole. Both AL- and DL-fed 3xTg female and male mice failed to locate the target hole within the given 90 seconds of assessment time (**Figs. 8A and 8F**). In contrast, CR-fed mice rapidly located the escape box during the training phase, as well as during the STM test and more profoundly during the LTM test conducted 12 days after the initial assay (**Figs. 8B and 8G**). Additionally, AL-fed and DL-fed 3xTg mice of both sexes made more errors during the training phases than CR-fed mice (**Figs. 8C-D, H-I**).

**Figure 8:**
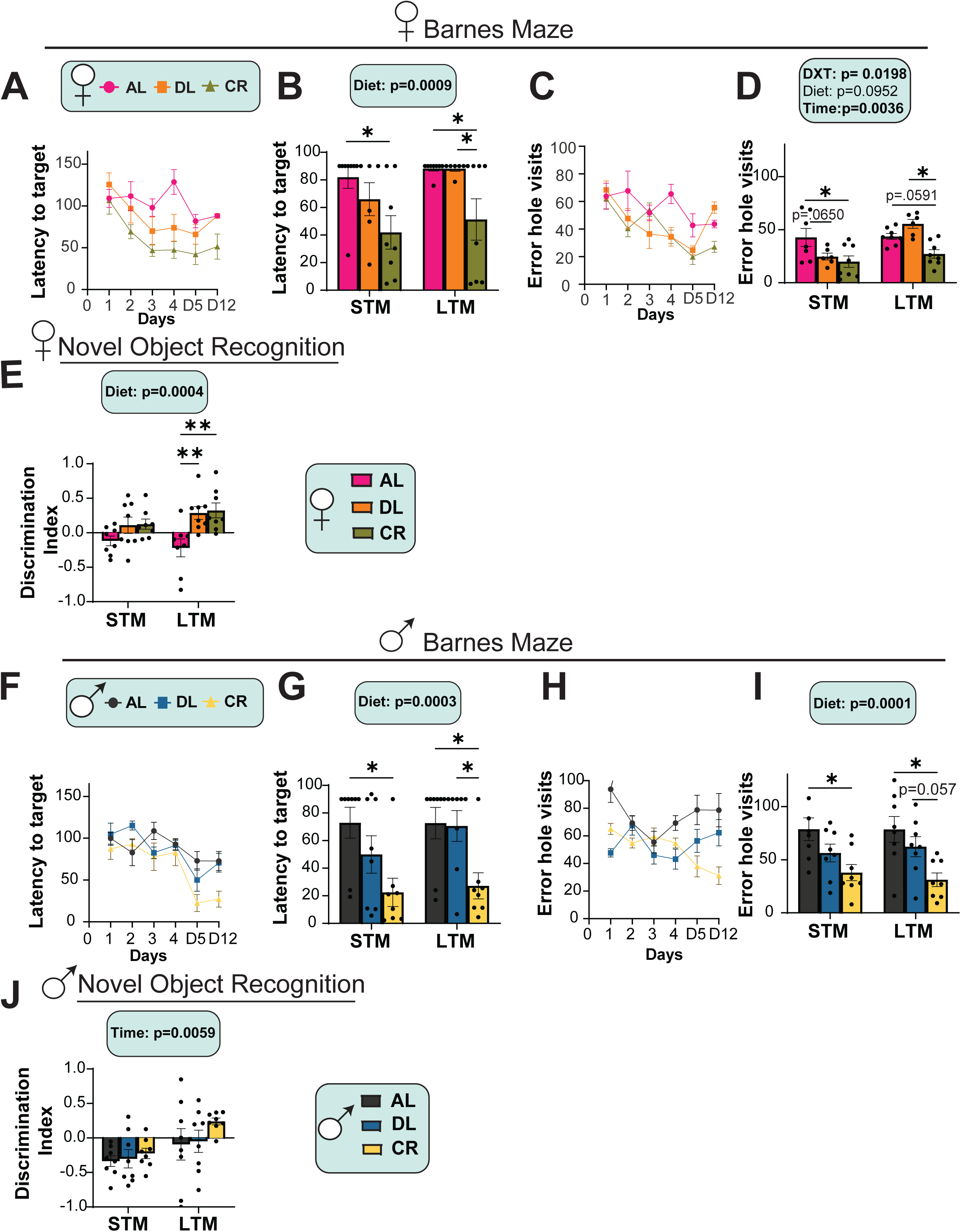
Fasting required for spatial memory recognition in both sexes. (A-J) The behavior of female and male 3xTg-AD mice was examined at 12 months of age after mice were fed the indicated diets for 6 months. (A-B) Latency of target in Barnes Maze acquisition period over the five days of training (A) and in short term memory (STM) and long-term memory (LTM) tests by female mice (B). (C-D) The number of error hole visits during Barnes maze training phase (C) and in STM and LTM tests by female mice (D). (E) The preference for a novel object over a familiar object was assayed in female mice via STM and LTM tests. (A-E) n=8 AL, n=6 DL and n=8 CR fed 3xTg biologically independent female mice. (F-G) Latency of target in Barnes Maze acquisition period over the five days of training (F) and in short term memory (STM) and long-term memory (LTM) tests by male mice (G). (H-I). The number of error hole visits during Barnes maze training phase (H) and in STM and LTM tests by male mice (I). (J) The preference for a novel object over a familiar object was assayed in male mice via STM and LTM tests. (F-J) n=8 AL, n=8 DL and n=8 CR fed 3xTg biologically independent male mice.(B,D,E,G,I,J) Statistics for the overall effects of diet, time and the interaction represent the p value from a 2-way ANOVA, *p<0.05, from a Sidak’s post-test examining the effect of parameters identified as significant in the 2-way ANOVA. Data represented as mean ± SEM.

The NOR task evaluates the preference for exploring a familiar object versus a new object and are quantified based on a discrimination index (DI). A positive DI implies a preference for exploring novelty, indicating that the memory of the familiar object persists and that the mice favor exploring the new object. We found that AL-fed 3xTg females spent more time with the familiar object than the novel object, suggesting defects in recognition memory (**Fig. 8E**). However, both the DL fed, and CR-fed females spent more time with novel objects than with the familiar object (**Fig. 8E**). However, 3xTg males did not show significant changes in their discrimination index (**Fig. 8J**). Overall, these results demonstrate that fasting plays a role in the cognitive benefits of a CR diet for 3xTg mice of both sexes.

### Fasting promotes survival outcomes in 3xTg AD-mice

Throughout our experiments, and consistent with our previous findings ^17^ as well as studies by other groups, we observed that 3xTg mice, particularly males, had a higher mortality rate as they approached one year of age. In agreement with these previous results, we found that AL-fed 3xTg males have a shorter lifespan than AL-fed female mice (**Figs. 9A-B**). 3xTg males fed a DL diet had significantly decreased survival (log-rank test, p=0.002) relative to both AL-fed and CR-fed males (**Fig. 9B**). In contrast, female 3xTg mice did not show significant differences in survival between the different dietary groups (**Fig. 9A**).

**Figure 9:**
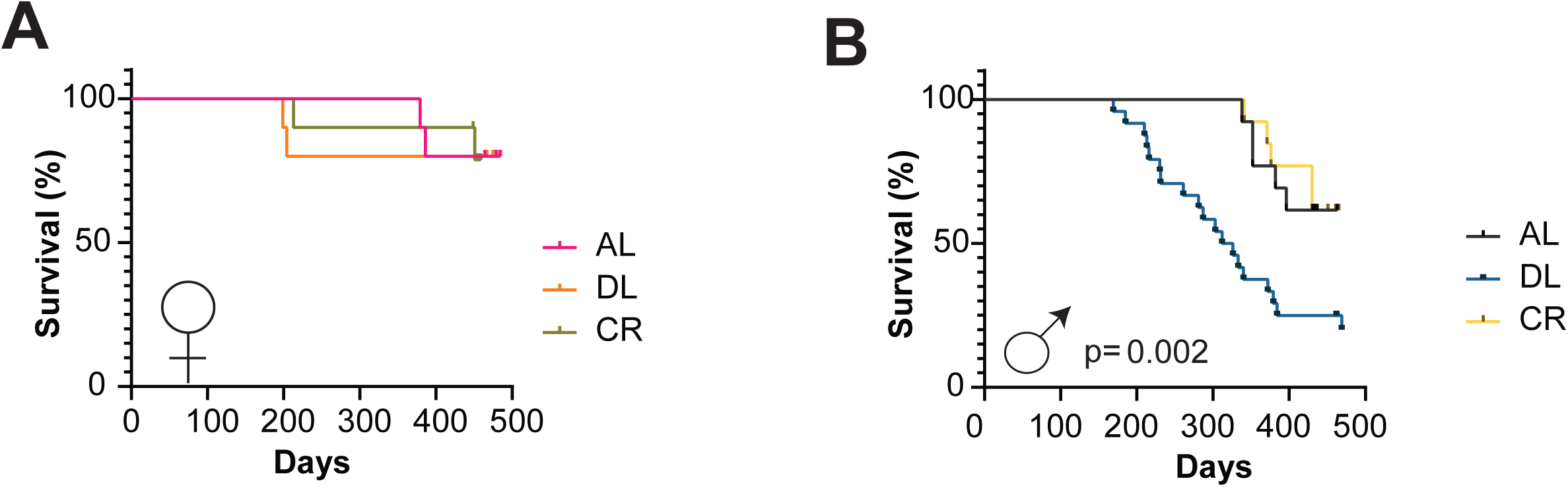
Fasting but not energy restriction alone affects the survival of male 3xTg mice. (A-B) Kaplan-Meier plots of the survival of female (A) and male (B) 3xTg female mice fed the indicated diets starting at 6 months of age. (A) For females: n=10 3xTg biologically independent mice/group (B) For males n=13 AL, n=23 DL and n=13 CR fed 3xTg biologically independent mice. p=0.002, log-rank test AL vs. DL mice.

## Discussion

Numerous studies have suggested that CR or intermittent fasting can promote cognitive function and protect from worsening AD pathology in animal models of AD ^4–8,32,33^. However, the physiological and molecular mechanisms that underlie the benefits of CR on cognition – as well as its beneficial impacts on the development and progression of Alzheimer’s disease – have remained elusive. Recent work from our lab and others has demonstrated that in wild-type mice, many of the benefits of CR arise not from the reduction in calories, but instead from the prolonged fasting period that accompanies once-a-day experimental CR regimen. Fasting in the context of CR is required for the benefits on insulin sensitivity, frailty, cognition, and lifespan in wild-type mice ^12,34^, and indeed fasting alone without reduction of calories is sufficient for many of the molecular effects of CR as well as extending lifespan in wild-type mice ^12,35^.

Here, we build on this foundation by determining if the protective effects of CR on the progression and development of AD arise from the restriction of energy alone or as a result of fasting between meals. We placed both female and male 3xTg AD mice on different feeding regimens allowing us to distinguish between the effects of energy restriction and fasting. We found that a reduction of calories without a prolonged fast improves glucose tolerance and aspects of AD pathology, reducing the density of Aβ plaques and neuroinflammation, but did not improve tau pathology. In contrast, prolonged fasting was necessary for CR-mediated improvements in insulin sensitivity and tau pathology. Without exception, CR with fasting had equal or greater cognitive benefits in multiple assays and in both sex of mice than energy restriction alone. Fasting was likewise required for CR-mediated molecular changes, including CR-induced reduction in brain mTOR signaling and the activation of autophagy. While there were sex-specific effects of CR, the importance of fasting largely held true across both sexes. Our results suggest that prolonged fasting between meals is an essential component of the benefits of CR on the development and progression of AD in mice.

CR-mediated attenuation of Aβ plaque deposition has been demonstrated in various AD mouse models ^5,7,36^. Further, a previous study, found that CR reduces both Aβ and tau accumulation and improved cognitive performance in the 3xTg mouse model of AD ^4^. Our findings are consistent with these previous reports, as we observed attenuation of Aβ plaques and tau pathology in CR-fed 3xTg mice.

Importantly, we have gained new insight into the mechanisms behind the benefits of a CR diet. First, DL-fed mice, which were energy restricted but did not experience prolonged daily fasts, showed significant attenuation of Aβ plaques. Notably, this occurred in the absence of improved insulin sensitivity, neuroinflammation, brain mTORC1 signaling, or activation of brain autophagy – all physiological and molecular changes that have been hypothesized to explain the ability of CR to slow the development or promote the clearance of Aβ plaques. Moreover, while DL-fed mice did show a trend towards cognitive improvements, it was clearly not as beneficial as CR. The modest benefits of targeting Aβ plaques alone may help explain the only partial success of Aβ-targeted antibodies in slowing cognitive decline in humans with AD ^37,38^.

In contrast, energy restriction with fasting – traditional once-per-day CR – was highly effective in improving cognition, reducing not only Aβ plaques but also hyperphosphorylation of tau and tau pathology, reducing neuroinflammation, and was accompanied by molecular changes believed to be beneficial, notably the reduction of mTORC1 signaling and activation of autophagy. In combination with our results in DL-fed mice, we conclude that prolonged fasting between meals is, at a minimum, required for most of the benefits of CR except the reduction of Aβ plaques. Rather surprisingly, this suggests that the benefits of reduced mTORC1 signaling, and increased autophagy may be mediated primarily by an Aβ-independent mechanism and moreover, previous studies have demonstrated that mTORC1 inhibition leads to the suppression of tauopathy ^39,40^. These findings are consistent with studies in transgenic mouse models of tauopathies, where CR has been shown to alleviate tau pathology and improve cognitive function ^41–44^. Intriguingly, despite reduced plaque deposition, we also observed increased astrocytic and microglial activation in the DL mice. However, this increase in glial activation was not observed in CR-fed females, suggesting that while caloric reduction alone may attenuate plaque pathology, the reduction in plaque deposition observed in the DL-fed mice may not be mediated by glial cells.

Throughout our study, we observed largely similar physiological, molecular and behavioral outcomes in response to all the three feeding regimens in both sexes with the exception in neuropathological outcomes. Specifically, the reduction of calories with or without fasting resulted in sex specific reduction of Aβ plaques in both cortex and hippocampus. While DL and CR reduced plaque deposition in both regions in females, there was no effect of either DL or CR on plaques in males.

As expected with CR, both males and females had reduced mTORC1 signaling and autophagy activation. However, females displayed significant reduction in mTORC1 when compared to DL-fed mice, whereas males showed significant reduction when compared to AL-fed mice. The extent of autophagy activation by CR was similar in both sexes, though induction of BDNF by CR was more pronounced in males following energy restriction.

These sex differences were even more clearly seen in our metabolomic analysis of plasma and brain tissue in the 3xTg mice. We observed significant changes in metabolic pathways in response to diet in females, with some overlap in the response to DL and CR feeding, while males showed many fewer changes with no overlapping changes between the DL and CR groups. One key pathway identified in females across both plasma and brain tissues was the cysteine and methionine metabolism pathway. This is particularly interesting given that methionine restriction has been previously shown to improve metabolic outcomes as well as improve longevity in animal models ^45–47^. Furthermore, components of the circulating methionine cycle are known to play an important role in the pathogenesis of AD ^48,49^. We observed changes in several methionine related metabolites which might suggest a potential role for epigenetic modifications in mediating some of the effects of CR and fasting, particularly in AD. Interestingly, we found a significant overlap in metabolite abundance between the DL and CR groups in the brains of female 3xTg mice. These overlapping metabolites were primarily amino acids, which were increased in both DL and CR groups compared to control mice. Consistent with previous studies in mice, we also observed an increase in branched-chain amino acids (Leucine, Isoleucine, and Valine) in the brains of both DL and CR-fed female mice ^12,50^.

The current study has several limitations that need to be addressed in future research. One significant limitation is the use of a diet diluted with indigestible cellulose (DL); while potentially a low-energy diet could be deleterious for a variety of reasons, in wild-type mice energy restriction induced by a low energy diet has similar impacts on lifespan as to energy restriction induced by many feeding small meals of a normal diet ^34^. We also did not study the effects of fasting alone, without energy restriction, on AD pathology and cognition in 3xTg mice. Based on the results we and others have observed in wild-type mice ^12,35^, we hypothesize that fasting may be sufficient to recapitulate many of the benefits of CR on cognition and AD pathology, but this has not been tested. Additionally, while there are many benefits of using the 3xTg mouse model of AD^53^, the use of other AD mouse models could provide additional insight, particularly into whether the benefits of CR are mediated via its remediations of tauopathy.

Our molecular analyses were limited; while we examined the effects of mTOR signaling and autophagy, we did not take a broader and unbiased view of the molecular pathways engaged by CR; fasting and energy restriction likely impinge on many other pathways which have been shown to impact AD pathology, e.g. AMPK and SIRT1 ^54–56^. Our molecular analysis did not explore the transcriptome or the proteome and was limited to the whole brain and plasma, rather than narrowing in on specific brain regions. A growing body of literature suggests other tissues outside the brain, including the liver, can also influence the development and progression of AD ^57–59^. Lastly, our study focused exclusively on the brain and plasma. Given the significant results we observed in insulin sensitivity and the overlap in metabolites in the brain, it would be important to include analyses of the liver, skeletal muscle, and adipose tissue in future studies. These tissues are key contributors to metabolic health, and examining their responses could provide important insights into the systemic effects of CR and fasting in AD mice.

In conclusion, our findings strongly suggest that while not all benefits of CR require fasting, fasting plays a critical role in CR-induced improvements in AD pathology. Although energy restriction alone was sufficient to mitigate plaque pathology in 3xTg females and improve certain aspects of cognition, fasting was essential for the reduction of tau pathology as well as many other molecular and histological markers of AD and the full cognitive benefits of CR. Our results suggest that CR with fasting – or, potentially, although we have not tested it here, fasting alone – could be a promising therapeutic approach to AD. Even if this diet regiment is too arduous for many or for the elderly in particular our results suggest that potential CR mimetic agents should aim to mimic the results of daily fasting, and not simply energy restriction, to full capture the benefits of CR for AD.

## Materials and Methods

### Animals

All procedures were performed in accordance with institutional guidelines and were approved by the Institutional Animal Care and Use Committee (IACUC) of the William S. Middleton Memorial Veterans Hospital and the University of Wisconsin-Madison IACUC (Madison, WI, USA). Male and female homozygous 3xTg-AD mice and their non-transgenic littermates were obtained from The Jackson Laboratory (Bar Harbor, ME, USA) and were bred and maintained at the vivarium with food and water available *ad libitum*. Prior to the start of the experiments at 6 months they were randomly assigned to different groups based on their body weight and diet. Mice were acclimatized on a 2018 Teklad Global 18% Protein Rodent Diet for 1 week before randomization and were singly housed. All mice were maintained at a temperature of approximately 22°C, and health checks were completed on all mice daily.

At the start of the experiment, mice were randomized to three different groups (1) AL, ad libitum diet; (2) DL, Diluted ad libitum diet where animals were provided with ad libitum access to a low-energy diet diluted with indigestible cellulose, which reduced caloric intake by ∼30% and (3) CR, animals in which calories were restricted by 30%, and animals were fed once per day during the start of the light period; Diet descriptions, compositions and item numbers are provided in **Table S1.**

### In vivo Procedures

Glucose tolerance test was performed by fasting the mice overnight and then injecting glucose (1 g kg^−1^) intraperitoneally (i.p.) as previously described ^47,60^. For insulin tolerance we fasted the mice for 4 hours and injected insulin intraperitoneally (0.75 U kg ^1^). Glucose measurements were taken using a Bayer Contour blood glucose meter (Bayer, Leverkusen, Germany) and test strips. Mouse body composition was determined using an EchoMRI Body Composition Analyzer (EchoMRI, Houston, TX, USA). For determining metabolic parameters [O2, CO2, food consumption, respiratory exchange ratio (RER), energy expenditure] and activity tracking, the mice were acclimated to housing in an Oxymax/CLAMS-HC metabolic chamber system (Columbus Instruments) for ∼24 h and data from a continuous 24 h period was then recorded and analyzed. Mice were euthanized by cervical dislocation after a 3 hr fast and tissues for molecular analysis were flash-frozen in liquid nitrogen or fixed and prepared as described in the methods below.

### Behavioral assays

All mice underwent behavioral phenotyping when they were twelve months old. The Novel object recognition test (NOR) was performed in an open field where the movements of the mouse were recorded via a camera that is mounted above the field. Before each test mice were acclimatized in the behavioral room for 30 minutes and were given a 5 min habituation trial with no objects on the field. This was followed by test phases that consisted of two trials that are 24 hrs apart: Short term memory test (STM and Long-term memory test (LTM). In the first trial, the mice were allowed to explore two identical objects placed diagonally on opposite corners of the field for 5 minutes. Following an hour after the acquisition phase, STM was performed and 24 hrs later, LTM was done by replacing one of the identical objects with a novel object. The results were quantified using a discrimination index (DI), representing the duration of exploration for the novel object compared to the old object.

For Barnes maze, the test involves 3 phases: habituation, acquisition training and the memory test. During habituation, mice were placed in the arena and allowed to freely explore the escape hole, escape box, and the adjacent area for 2 min. Following that during acquisition training the mice were given 180s to find the escape hole, and if they failed to enter the escape box within that time, they were led to the escape hole. After 4 days of training, on the 5^th^ day (STM) and 12^th^ day (LTM) the mice were given 90s memory probe trials. The latency to enter the escape hole, distance traveled, and average speed were analyzed using Ethovision XT (Noldus).

### Immunoblotting

Tissue samples from brain were lysed in cold RIPA buffer supplemented with phosphatase inhibitor and protease inhibitor cocktail tablets (Thermo Fisher Scientific, Waltham, MA, USA) using a FastPrep 24 (M.P. Biomedicals, Santa Ana, CA, USA) with bead-beating tubes (16466– 042) from (VWR, Radnor, PA, USA) and zirconium ceramic oxide bulk beads (15340159) from (Thermo Fisher Scientific, Waltham, MA, USA). Protein lysates were then centrifuged at 13,300 rpm for 10 min and the supernatant was collected. Protein concentration was determined by Bradford (Pierce Biotechnology, Waltham, MA, USA). 20 μg protein was separated by SDS– PAGE (sodium dodecyl sulfate–polyacrylamide gel electrophoresis) on 8%, 10%, or 16% resolving gels (ThermoFisher Scientific, Waltham, MA, USA) and transferred to PVDF membrane (EMD Millipore, Burlington, MA, USA). The phosphorylation status of mTORC1 substrates including p-S240/S244 S6 and 4E-BP1 T37/S46 were assessed in the brain along with autophagy receptor p62 (sequestosome 1, SQSTM1) and the autophagosome marker LC3A/B. Tau pathology was assessed by western blot with anti-tau antibody. Antibody vendors, catalog numbers and the dilution used are provided in **Table S12.** Imaging was performed using a Bio-Rad Chemidoc MP imaging station (Bio-Rad, Hercules, CA, USA). Quantification was performed by densitometry using NIH ImageJ software.

### Histology for AD neuropathology markers

Mice were euthanized by cervical dislocation after a 3 hour fast, and the right hemisphere was fixed in formalin for histology whereas the left hemisphere was snap-frozen for biochemical analysis. For amyloid plaque staining, briefly brain sections were deparaffinized and rehydrated according to standard protocol. For epitope retrieval, mounted slides were pretreated in 70% formic acid at room temperature for 10 min. Tissue sections were subsequently blocked with normal goat serum (NGS) at room temperature for 1 hr, then incubated with monoclonal antibodies 6E10 (1:100), at 4°C overnight. Aβ immunostained profiles were visualized using diaminobenzidine chromagen. For p-Tau staining and glial activation, brains were analyzed with anti-GFAP (astrocytic marker), and anti-Iba1 (microglial marker) antibodies respectively. The following primary antibodies were used: phospho-Tau (Thr231) monoclonal antibody (AT180) (Thermo Fisher Scientific; # MN1040, 1:100) anti-GFAP (Thermo Fisher; # PIMA512023; 1:1,000), anti-IBA1 (Abcam; #ab178847; 1:1,000). Sections were imaged using an EVOS microscope (Thermo Fisher Scientific Inc., Waltham, MA, USA) at a magnification of 4X, 10X and 40X magnification. Image-J was used for quantification by converting images into binary images via an intensity threshold and positive area was quantified. ^61^.

### Targeted Metabolomics on Plasma

#### Metabolite extraction

Plasma (20μL of) was transferred to an individual 1.5 mL microcentrifuge tube and incubated with 400μL −80°C 80:20 Methanol (MeOH): H2O extraction solvent on dry ice for 5 minutes post-vortexing. Serum homogenate was centrifuged at 21,000 xg for 5 minutes at 4°C. Supernatant was transferred to 1.5 mL microcentrifuge tube after which the remining pellet was resuspended in 400 µL −20°C 40:40:20 Acetonitrile (ACN): MeOH: H2O extraction solvent and incubated on ice for 5minutes. Serum homogenate was again centrifuged at 21,000 xg for 5 minutes at 4°C after which the supernatant was pooled with the previously isolated metabolite fraction. The 40:20:20 ACN: MeOH:H_2_O extraction was then repeated as previously described. Next, the pooled metabolite extract for each sample was transferred to a 1.5 mL microcentrifuge Eppendorf tube and completely dried using a Thermo Fisher Savant ISS110 SpeedVac. Dried metabolite extracts were resuspended in 100 µL of LCMS-graded water following microcentrifugation for 5 minutes at 21,000 xg at 4C to pellet any remaining insoluble debris. Supernatant was then transferred to a glass vial for LC-MS analysis.

### Targeted Metabolomics on Brain

#### Metabolite Extraction

Approximately 10mg of tissue was homogenized on ice using a dounce homogenizer in 400µL of 80:20 methanol. The homogenate was then transferred to a 1.5mL microcentrifuge tube. An additional 100µL of the solvent was used to rinse the mortar and was subsequently added to the same tube. The mixture was incubated on ice for 5 minutes before being centrifuged at 21,000 xg for 5 minutes at 4°C. After centrifugation, 500µL of the supernatant was transferred to a fresh 1.5mL microcentrifuge tube and dried completely using a Thermo Fisher Savant ISS110 SpeedVac. The protein pellet was resuspended in 100µL RIPA buffer (10mM Tris-HCl, pH 8.0, 1mM EDTA, 0.5mM EGTA, 1% Triton X-100, 0.1% Sodium Deoxycholate, 0.1% SDS, and 140mM NaCl) to normalize the intensity values for each targeted metabolite. The dried metabolite extracts were resuspended in 100L of LC-MS-grade water and centrifuged again at 21,000 xg for 5 minutes at 4 °C to pellet any remaining insoluble debris. The resulting supernatant was then transferred to a glass vial for LCMS analysis.

### LC-MS Metabolite Analysis

Each prepared metabolite sample was injected onto a Thermo Fisher Scientific Vanquish UHPLC with a Waters Acquity UPLC BEH C18 column (1.7μm, 2.1×100mm; Waters Corp., Milford, MA, USA) and analyzed using a Thermo Fisher Q Exactive orbitrap mass spectrometer in negative ionization mode. LC separation was performed over a 25 minute method with a 14.5 minute linear gradient of mobile phase (buffer A, 97% water with 3% methanol, 10mM tributylamine, and acetic acid-adjusted pH of 8.3) and organic phase (buffer B, 100% methanol) (0 minute, 5% B; 2.5 minute, 5% B; 17 minute, 95% B; 19.5 minute, 5% B; 20 minute, 5% B; 25 minute, 5% B, flow rate 0.2mL/min). A quantity of 10µL of each sample was injected into the system for analysis. The ESI settings were 30/10/1 for sheath/aux/sweep gas flow rates, 2.50kV for spray voltage, 50 for S-lens RF level, 350C for capillary temperature, and 300C for auxiliary gas heater temperature. MS1 scans were operated at resolution = 70,000, scan range = 85-1250m/z, automatic gain control target = 1 x 10^6^, and 100ms maximum IT. Metabolites were identified and quantified using El-MAVEN (v0.12.1-beta) with metabolite retention times empirically determined in-house. Peak Area top values were imported into MetaboAnalyst for statistical analysis (one factor) using default settings.

### Metabolomics analysis

The metabolites were initially normalized using a log base 2 transformation. Additionally, to control for false positives, P values were adjusted using the Benjamini–Hochberg procedure with a false discovery rate (FDR) of 20%. Subsequently, pathway analysis was conducted using the online tool, El-MAVEN (https://docs.polly.elucidata.io/Apps/Metabolomic%20Data/El-MAVEN.html). The Pathway Analysis function of the tool was employed by inputting a list of significantly altered metabolites based on their corresponding human metabolome database (HMDB) IDs obtained from the linear model with a significance threshold of p<0.05.

### Statistical Analysis

All statistical analyses were conducted using Prism, version 9 (GraphPad Software Inc., San Diego, CA, USA). Tests involving multiple factors were analyzed by either a two-way analysis of variance (ANOVA) with diet and time or sex as variables or by one-way ANOVA, followed by a Dunnett’s, Tukey-Kramer, or Sidak’s post-hoc test as specified in the figure legends. Metabolomics data were analysed using metaboanalyst (version 6.0). Kaplan–Meir survival analysis of 3xTg mice was performed with log-rank comparisons stratified by sex and diet. Cox proportional hazards analysis of 3xTg mice was performed using sex and diet as covariates. Alpha was set at 5% (p < .05 considered to be significant). Data are presented as the mean ± SEM unless otherwise specified.

## ACKNOWLEDGEMENTS

We would like to thank Dr. Heidi Pak for her valuable insights and support. We thank all the members of the Lamming lab for their feedback. The Lamming lab is supported in part by the NIA (AG056771, AG062328, AG061635, AG081482, and AG084156), the NIDDK (DK125859), by a grant from the Alzheimer’s Association (23AARG-1029665), and by startup funds from UW-Madison. RB was supported in part by F31AG081115. MMS was supported in part by a Supplement to Promote Diversity in Health-Related Research RF1AG056771-06S1. CLG was supported in part by Dalio Philanthropies, a Glenn Foundation for Medical Research Postdoctoral Fellowship, and by grant HF-AGE AGE-009 from the Hevolution Foundation to CLG. MFC was supported in part by F31 AG082504. The Puglielli lab is supported in part by the NINDS (NS094154), the NIGMS (GM148487) and the NIA (AG078794). The Denu lab is supported in part by the NIH (GM149279 and DK125859). The authors thank the University of Wisconsin Carbone Cancer Center Experimental Animal Pathology Laboratory supported by P30 CA014520, for use of its facilities and services. The Lamming lab was supported in part by the U.S. Department of Veterans Affairs (I01-BX004031 and IS1-BX005524), and this work was supported using facilities and resources from the William S. Middleton Memorial Veterans Hospital. The content is solely the responsibility of the authors and does not necessarily represent the official views of the NIH. This work does not represent the views of the Department of Veterans Affairs or the United States Government.

## DATA AVAILABILITY

The authors declare that source data supporting the findings of this study are available within the paper and its supplementary information and Source Data files. Targeted brain metabolomics data has been deposited in MassIVE with accession code MSV000095847 [doi:10.25345/C5N873B30]. Targeted plasma metabolomics data has been deposited in MassIVE with accession code MSV000095887 [doi:10.25345/C5GF0N767].

## AUTHOR CONTRIBUTIONS

RB, JHH, MMS, CLG, RM, AT, CYY, MFC, SS, JI, IG, and YL conducted the experiments. RB, JHH, CLG, DV and DWL analyzed the data. RB, JHH, MJB, DH, JMD, LP and DWL wrote and edited the manuscript.

## References

1. Rajan, K.B., Weuve, J., Barnes, L.L., McAninch, E.A., Wilson, R.S., and Evans, D.A. (2021). Population estimate of people with clinical Alzheimer’s disease and mild cognitive impairment in the United States (2020-2060). Alzheimer’s & dementia : the journal of the Alzheimer’s Association 17, 1966–1975. 10.1002/alz.12362.

2. 2023 Alzheimer’s disease facts and figures. (2023). Alzheimer’s & dementia : the journal of the Alzheimer’s Association 19, 1598–1695. 10.1002/alz.13016.

3. Green, C.L., Lamming, D.W., and Fontana, L. (2022). Molecular mechanisms of dietary restriction promoting health and longevity. Nat Rev Mol Cell Biol 23, 56–73. 10.1038/s41580-021-00411-4.

4. Halagappa, V.K., Guo, Z., Pearson, M., Matsuoka, Y., Cutler, R.G., Laferla, F.M., and Mattson, M.P. (2007). Intermittent fasting and caloric restriction ameliorate age-related behavioral deficits in the triple-transgenic mouse model of Alzheimer’s disease. Neurobiol Dis 26, 212–220. 10.1016/j.nbd.2006.12.019.

5. Mouton, P.R., Chachich, M.E., Quigley, C., Spangler, E., and Ingram, D.K. (2009). Caloric restriction attenuates amyloid deposition in middle-aged dtg APP/PS1 mice. Neurosci Lett 464, 184–187. 10.1016/j.neulet.2009.08.038.

6. Schafer, M.J., Alldred, M.J., Lee, S.H., Calhoun, M.E., Petkova, E., Mathews, P.M., and Ginsberg, S.D. (2015). Reduction of beta-amyloid and gamma-secretase by calorie restriction in female Tg2576 mice. Neurobiol Aging 36, 1293–1302. 10.1016/j.neurobiolaging.2014.10.043.

7. Wang, J., Ho, L., Qin, W., Rocher, A.B., Seror, I., Humala, N., Maniar, K., Dolios, G., Wang, R., Hof, P.R., and Pasinetti, G.M. (2005). Caloric restriction attenuates beta-amyloid neuropathology in a mouse model of Alzheimer’s disease. FASEB J 19, 659–661. 10.1096/fj.04-3182fje.

8. Qin, W., Chachich, M., Lane, M., Roth, G., Bryant, M., de Cabo, R., Ottinger, M.A., Mattison, J., Ingram, D., Gandy, S., and Pasinetti, G.M. (2006). Calorie restriction attenuates Alzheimer’s disease type brain amyloidosis in Squirrel monkeys (Saimiri sciureus). Journal of Alzheimer’s disease : JAD 10, 417–422. 10.3233/jad-2006-10411.

9. Witte, A.V., Fobker, M., Gellner, R., Knecht, S., and Floel, A. (2009). Caloric restriction improves memory in elderly humans. Proc Natl Acad Sci U S A 106, 1255–1260. 10.1073/pnas.0808587106.

10. Acosta-Rodriguez, V.A., de Groot, M.H.M., Rijo-Ferreira, F., Green, C.B., and Takahashi, J.S. (2017). Mice under Caloric Restriction Self-Impose a Temporal Restriction of Food Intake as Revealed by an Automated Feeder System. Cell Metab 26, 267–277 e262. 10.1016/j.cmet.2017.06.007.

11. Wahl, D., Coogan, S.C., Solon-Biet, S.M., de Cabo, R., Haran, J.B., Raubenheimer, D., Cogger, V.C., Mattson, M.P., Simpson, S.J., and Le Couteur, D.G. (2017). Cognitive and behavioral evaluation of nutritional interventions in rodent models of brain aging and dementia. Clin Interv Aging 12, 1419–1428. 10.2147/CIA.S145247.

12. Pak, H.H., Haws, S.A., Green, C.L., Koller, M., Lavarias, M.T., Richardson, N.E., Yang, S.E., Dumas, S.N., Sonsalla, M., Bray, L., et al. (2021). Fasting drives the metabolic, molecular and geroprotective effects of a calorie-restricted diet in mice. Nat Metab 3, 1327–1341. 10.1038/s42255-021-00466-9.

13. Oddo, S., Caccamo, A., Shepherd, J.D., Murphy, M.P., Golde, T.E., Kayed, R., Metherate, R., Mattson, M.P., Akbari, Y., and LaFerla, F.M. (2003). Triple-transgenic model of Alzheimer’s disease with plaques and tangles: intracellular Abeta and synaptic dysfunction. Neuron 39, 409–421. 10.1016/s0896-6273(03)00434-3.

14. Bruss, M.D., Khambatta, C.F., Ruby, M.A., Aggarwal, I., and Hellerstein, M.K. (2010). Calorie restriction increases fatty acid synthesis and whole body fat oxidation rates. Am J Physiol Endocrinol Metab 298, E108–116. 10.1152/ajpendo.00524.2009.

15. Ansoleaga, B., Jove, M., Schluter, A., Garcia-Esparcia, P., Moreno, J., Pujol, A., Pamplona, R., Portero-Otin, M., and Ferrer, I. (2015). Deregulation of purine metabolism in Alzheimer’s disease. Neurobiol Aging 36, 68–80. 10.1016/j.neurobiolaging.2014.08.004.

16. Palmer, A.M. (1999). The activity of the pentose phosphate pathway is increased in response to oxidative stress in Alzheimer’s disease. J Neural Transm (Vienna) 106, 317–328. 10.1007/s007020050161.

17. Babygirija, R., Sonsalla, M.M., Mill, J., James, I., Han, J.H., Green, C.L., Calubag, M.F., Wade, G., Tobon, A., Michael, J., et al. (2024). Protein restriction slows the development and progression of pathology in a mouse model of Alzheimer’s disease. Nature communications 15, 5217. 10.1038/s41467-024-49589-z.

18. Talboom, J.S., Velazquez, R., and Oddo, S. (2015). The mammalian target of rapamycin at the crossroad between cognitive aging and Alzheimer’s disease. NPJ Aging Mech Dis 1, 15008. 10.1038/npjamd.2015.8.

19. Uddin, M.S., Mamun, A.A., Labu, Z.K., Hidalgo-Lanussa, O., Barreto, G.E., and Ashraf, G.M. (2019). Autophagic dysfunction in Alzheimer’s disease: Cellular and molecular mechanistic approaches to halt Alzheimer’s pathogenesis. J Cell Physiol 234, 8094–8112. 10.1002/jcp.27588.

20. Simcox, J., and Lamming, D.W. (2022). The central moTOR of metabolism. Dev Cell 57, 691–706. 10.1016/j.devcel.2022.02.024.

21. Duan, W., Lee, J., Guo, Z., and Mattson, M.P. (2001). Dietary restriction stimulates BDNF production in the brain and thereby protects neurons against excitotoxic injury. J Mol Neurosci 16, 1–12. 10.1385/JMN:16:1:1.

22. Kishi, T., Hirooka, Y., Nagayama, T., Isegawa, K., Katsuki, M., Takesue, K., and Sunagawa, K. (2015). Calorie restriction improves cognitive decline via up-regulation of brain-derived neurotrophic factor: tropomyosin-related kinase B in hippocampus ofobesity-induced hypertensive rats. Int Heart J 56, 110–115. 10.1536/ihj.14-168.

23. Miranda, M., Morici, J.F., Zanoni, M.B., and Bekinschtein, P. (2019). Brain-Derived Neurotrophic Factor: A Key Molecule for Memory in the Healthy and the Pathological Brain. Front Cell Neurosci 13, 363. 10.3389/fncel.2019.00363.

24. Muller, L., Power Guerra, N., Stenzel, J., Ruhlmann, C., Lindner, T., Krause, B.J., Vollmar, B., Teipel, S., and Kuhla, A. (2021). Long-Term Caloric Restriction Attenuates beta-Amyloid Neuropathology and Is Accompanied by Autophagy in APPswe/PS1delta9 Mice. Nutrients 13. 10.3390/nu13030985.

25. Phillips, H.S., Hains, J.M., Armanini, M., Laramee, G.R., Johnson, S.A., and Winslow, J.W. (1991). BDNF mRNA is decreased in the hippocampus of individuals with Alzheimer’s disease. Neuron 7, 695–702. 10.1016/0896-6273(91)90273-3.

26. Connor, B., Young, D., Yan, Q., Faull, R.L., Synek, B., and Dragunow, M. (1997). Brain-derived neurotrophic factor is reduced in Alzheimer’s disease. Brain Res Mol Brain Res 49, 71–81. 10.1016/s0169-328x(97)00125-3.

27. Hock, C., Heese, K., Hulette, C., Rosenberg, C., and Otten, U. (2000). Region-specific neurotrophin imbalances in Alzheimer disease: decreased levels of brain-derived neurotrophic factor and increased levels of nerve growth factor in hippocampus and cortical areas. Archives of neurology 57, 846–851. 10.1001/archneur.57.6.846.

28. Peng, S., Garzon, D.J., Marchese, M., Klein, W., Ginsberg, S.D., Francis, B.M., Mount, H.T., Mufson, E.J., Salehi, A., and Fahnestock, M. (2009). Decreased brain-derived neurotrophic factor depends on amyloid aggregation state in transgenic mouse models of Alzheimer’s disease. J Neurosci 29, 9321–9329. 10.1523/JNEUROSCI.4736-08.2009.

29. Ng, T.K.S., Ho, C.S.H., Tam, W.W.S., Kua, E.H., and Ho, R.C. (2019). Decreased Serum Brain-Derived Neurotrophic Factor (BDNF) Levels in Patients with Alzheimer’s Disease (AD): A Systematic Review and Meta-Analysis. Int J Mol Sci 20. 10.3390/ijms20020257.

30. Takei, N., Inamura, N., Kawamura, M., Namba, H., Hara, K., Yonezawa, K., and Nawa, H. (2004). Brain-derived neurotrophic factor induces mammalian target of rapamycin-dependent local activation of translation machinery and protein synthesis in neuronal dendrites. J Neurosci 24, 9760–9769. 10.1523/JNEUROSCI.1427-04.2004.

31. Takei, N., Kawamura, M., Hara, K., Yonezawa, K., and Nawa, H. (2001). Brain-derived neurotrophic factor enhances neuronal translation by activating multiple initiation processes: comparison with the effects of insulin. J Biol Chem 276, 42818–42825. 10.1074/jbc.M103237200.

32. Hu, Y., Yang, Y., Zhang, M., Deng, M., and Zhang, J.J. (2017). Intermittent Fasting Pretreatment Prevents Cognitive Impairment in a Rat Model of Chronic Cerebral Hypoperfusion. The Journal of nutrition 147, 1437–1445. 10.3945/jn.116.245613.

33. Liu, Y., Cheng, A., Li, Y.J., Yang, Y., Kishimoto, Y., Zhang, S., Wang, Y., Wan, R., Raefsky, S.M., Lu, D., et al. (2019). SIRT3 mediates hippocampal synaptic adaptations to intermittent fasting and ameliorates deficits in APP mutant mice. Nature communications 10, 1886. 10.1038/s41467-019-09897-1.

34. Acosta-Rodriguez, V., Rijo-Ferreira, F., Izumo, M., Xu, P., Wight-Carter, M., Green, C.B., and Takahashi, J.S. (2022). Circadian alignment of early onset caloric restriction promotes longevity in male C57BL/6J mice. Science 376, 1192–1202. 10.1126/science.abk0297.

35. Aon, M.A., Bernier, M., Mitchell, S.J., Di Germanio, C., Mattison, J.A., Ehrlich, M.R., Colman, R.J., Anderson, R.M., and de Cabo, R. (2020). Untangling Determinants of Enhanced Health and Lifespan through a Multi-omics Approach in Mice. Cell Metab 32, 100–116 e104. 10.1016/j.cmet.2020.04.018.

36. Patel, N.V., Gordon, M.N., Connor, K.E., Good, R.A., Engelman, R.W., Mason, J., Morgan, D.G., Morgan, T.E., and Finch, C.E. (2005). Caloric restriction attenuates Abeta-deposition in Alzheimer transgenic models. Neurobiol Aging 26, 995–1000. 10.1016/j.neurobiolaging.2004.09.014.

37. Mehta, D., Jackson, R., Paul, G., Shi, J., and Sabbagh, M. (2017). Why do trials for Alzheimer’s disease drugs keep failing? A discontinued drug perspective for 2010-2015. Expert Opin Investig Drugs 26, 735–739. 10.1080/13543784.2017.1323868.

38. Zhang, Y., Chen, H., Li, R., Sterling, K., and Song, W. (2023). Amyloid beta-based therapy for Alzheimer’s disease: challenges, successes and future. Signal Transduct Target Ther 8, 248. 10.1038/s41392-023-01484-7.

39. Tramutola, A., Lanzillotta, C., and Di Domenico, F. (2017). Targeting mTOR to reduce Alzheimer-related cognitive decline: from current hits to future therapies. Expert Rev Neurother 17, 33–45. 10.1080/14737175.2017.1244482.

40. Wilkinson, J.E., Burmeister, L., Brooks, S.V., Chan, C.C., Friedline, S., Harrison, D.E., Hejtmancik, J.F., Nadon, N., Strong, R., Wood, L.K., et al. (2012). Rapamycin slows aging in mice. Aging Cell 11, 675–682. 10.1111/j.1474-9726.2012.00832.x.

41. Cogut, V., McNeely, T.L., Bussian, T.J., Graves, S.I., and Baker, D.J. (2024). Caloric Restriction Improves Spatial Learning Deficits in Tau Mice. Journal of Alzheimer’s disease : JAD 98, 925–940. 10.3233/JAD-231117.

42. Wu, P., Shen, Q., Dong, S., Xu, Z., Tsien, J.Z., and Hu, Y. (2008). Calorie restriction ameliorates neurodegenerative phenotypes in forebrain-specific presenilin-1 and presenilin-2 double knockout mice. Neurobiol Aging 29, 1502–1511. 10.1016/j.neurobiolaging.2007.03.028.

43. Bejanin, A., Schonhaut, D.R., La Joie, R., Kramer, J.H., Baker, S.L., Sosa, N., Ayakta, N., Cantwell, A., Janabi, M., Lauriola, M., et al. (2017). Tau pathology and neurodegeneration contribute to cognitive impairment in Alzheimer’s disease. Brain 140, 3286–3300. 10.1093/brain/awx243.

44. Brownlow, M.L., Joly-Amado, A., Azam, S., Elza, M., Selenica, M.L., Pappas, C., Small, B., Engelman, R., Gordon, M.N., and Morgan, D. (2014). Partial rescue of memory deficits induced by calorie restriction in a mouse model of tau deposition. Behav Brain Res 271, 79–88. 10.1016/j.bbr.2014.06.001.

45. Wang, L., Ren, B., Zhang, Q., Chu, C., Zhao, Z., Wu, J., Zhao, W., Liu, Z., and Liu, X. (2020). Methionine restriction alleviates high-fat diet-induced obesity: Involvement of diurnal metabolism of lipids and bile acids. Biochim Biophys Acta Mol Basis Dis 1866, 165908. 10.1016/j.bbadis.2020.165908.

46. Gao, X., Sanderson, S.M., Dai, Z., Reid, M.A., Cooper, D.E., Lu, M., Richie, J.P., Jr., Ciccarella, A., Calcagnotto, A., Mikhael, P.G., et al. (2019). Dietary methionine influences therapy in mouse cancer models and alters human metabolism. Nature 572, 397–401. 10.1038/s41586-019-1437-3.

47. Yu, D., Yang, S.E., Miller, B.R., Wisinski, J.A., Sherman, D.S., Brinkman, J.A., Tomasiewicz, J.L., Cummings, N.E., Kimple, M.E., Cryns, V.L., and Lamming, D.W. (2018). Short-term methionine deprivation improves metabolic health via sexually dimorphic, mTORC1-independent mechanisms. FASEB J 32, 3471–3482. 10.1096/fj.201701211R.

48. Zhao, Y., Dong, X., Chen, B., Zhang, Y., Meng, S., Guo, F., Guo, X., Zhu, J., Wang, H., Cui, H., and Li, S. (2022). Blood levels of circulating methionine components in Alzheimer’s disease and mild cognitive impairment: A systematic review and meta-analysis. Front Aging Neurosci 14, 934070. 10.3389/fnagi.2022.934070.

49. Xi, Y., Zhang, Y., Zhou, Y., Liu, Q., Chen, X., Liu, X., Grune, T., Shi, L., Hou, M., and Liu, Z. (2023). Effects of methionine intake on cognitive function in mild cognitive impairment patients and APP/PS1 Alzheimer’s Disease model mice: Role of the cystathionine-beta-synthase/H(2)S pathway. Redox Biol 59, 102595. 10.1016/j.redox.2022.102595.

50. Collet, T.H., Sonoyama, T., Henning, E., Keogh, J.M., Ingram, B., Kelway, S., Guo, L., and Farooqi, I.S. (2017). A Metabolomic Signature of Acute Caloric Restriction. J Clin Endocrinol Metab 102, 4486–4495. 10.1210/jc.2017-01020.

51. Belfiore, R., Rodin, A., Ferreira, E., Velazquez, R., Branca, C., Caccamo, A., and Oddo, S. (2019). Temporal and regional progression of Alzheimer’s disease-like pathology in 3xTg-AD mice. Aging Cell 18, e12873. 10.1111/acel.12873.

52. Kane, A.E., Shin, S., Wong, A.A., Fertan, E., Faustova, N.S., Howlett, S.E., and Brown, R.E. (2018). Sex Differences in Healthspan Predict Lifespan in the 3xTg-AD Mouse Model of Alzheimer’s Disease. Front Aging Neurosci 10, 172. 10.3389/fnagi.2018.00172.

53. Sonsalla, M.M., and Lamming, D.W. (2023). Geroprotective interventions in the 3xTg mouse model of Alzheimer’s disease. Geroscience 45, 1343–1381. 10.1007/s11357-023-00782-w.

54. Ruhlmann, C., Wolk, T., Blumel, T., Stahn, L., Vollmar, B., and Kuhla, A. (2016). Long-term caloric restriction in ApoE-deficient mice results in neuroprotection via Fgf21-induced AMPK/mTOR pathway. Aging (Albany NY) 8, 2777–2789. 10.18632/aging.101086.

55. Cohen, H.Y., Miller, C., Bitterman, K.J., Wall, N.R., Hekking, B., Kessler, B., Howitz, K.T., Gorospe, M., de Cabo, R., and Sinclair, D.A. (2004). Calorie restriction promotes mammalian cell survival by inducing the SIRT1 deacetylase. Science 305, 390–392. 10.1126/science.1099196.

56. Madeo, F., Carmona-Gutierrez, D., Hofer, S.J., and Kroemer, G. (2019). Caloric Restriction Mimetics against Age-Associated Disease: Targets, Mechanisms, and Therapeutic Potential. Cell Metab 29, 592–610. 10.1016/j.cmet.2019.01.018.

57. Kaur, H., Seeger, D., Golovko, S., Golovko, M., and Combs, C.K. (2021). Liver Bile Acid Changes in Mouse Models of Alzheimer’s Disease. Int J Mol Sci 22. 10.3390/ijms22147451.

58. Bosoi, C.R., Vandal, M., Tournissac, M., Leclerc, M., Fanet, H., Mitchell, P.L., Verreault, M., Trottier, J., Virgili, J., Tremblay, C., et al. (2021). High-Fat Diet Modulates Hepatic Amyloid beta and Cerebrosterol Metabolism in the Triple Transgenic Mouse Model of Alzheimer’s Disease. Hepatol Commun 5, 446–460. 10.1002/hep4.1609.

59. Bassendine, M.F., Taylor-Robinson, S.D., Fertleman, M., Khan, M., and Neely, D. (2020). Is Alzheimer’s Disease a Liver Disease of the Brain? Journal of Alzheimer’s disease : JAD 75, 1–14. 10.3233/JAD-190848.

60. Bellantuono, I., de Cabo, R., Ehninger, D., Di Germanio, C., Lawrie, A., Miller, J., Mitchell, S.J., Navas-Enamorado, I., Potter, P.K., Tchkonia, T., et al. (2020). A toolbox for the longitudinal assessment of healthspan in aging mice. Nature protocols 15, 540–574. 10.1038/s41596-019-0256-1.

61. Rigby, M.J., Lawton, A.J., Kaur, G., Banduseela, V.C., Kamm, W.E., Lakkaraju, A., Denu, J.M., and Puglielli, L. (2021). Endoplasmic reticulum acetyltransferases Atase1 and Atase2 differentially regulate reticulophagy, macroautophagy and cellular acetyl-CoA metabolism. Commun Biol 4, 454. 10.1038/s42003-021-01992-8.

