## Supplemental Figures and Table Legends for "Fasting is required for many of the benefits of calorie restriction in the 3xTg mouse model of Alzheimer’s disease"

### Supplementary Fig 1

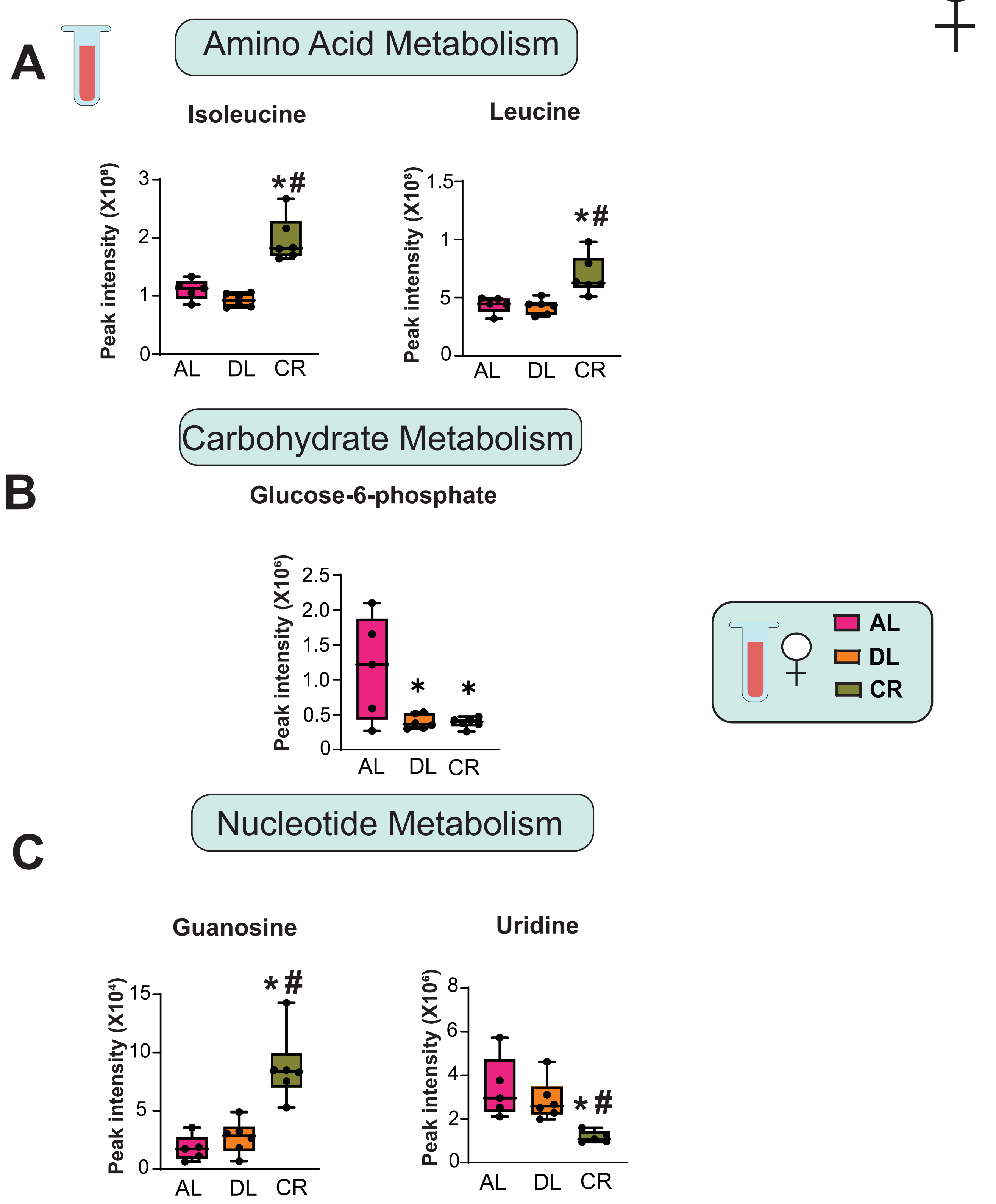

**Supplementary Figure 1: Relative abundance of significant plasma metabolites in female 3xTg-AD mice.**

(A-C) Targeted metabolomics were performed on the plasma of female 3xTg mice fed AL, Diluted AL and CR diets (n = 6) biologically independent mice per diet. (A) Relative abundance of significant amino acid metabolites (B) nucleotide metabolites and (C) carbohydrate metabolites. (A-C) * symbol represents a significant difference versus AL mice ($p \leq 0.05$); # symbol represents a significant difference versus Diluted AL mice ($p \leq 0.05$) based on Tukey's test post one-way ANOVA. Overlaid box plots show center as median and 25th-75thpercentiles; whiskers represent minima and maxima.

### Supplementary Figure 2

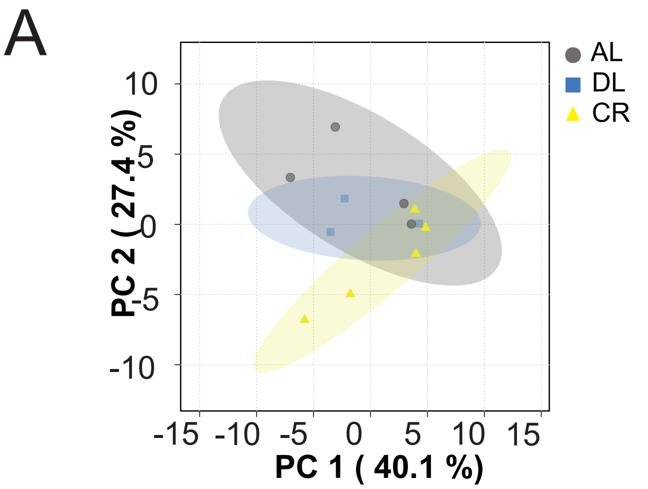


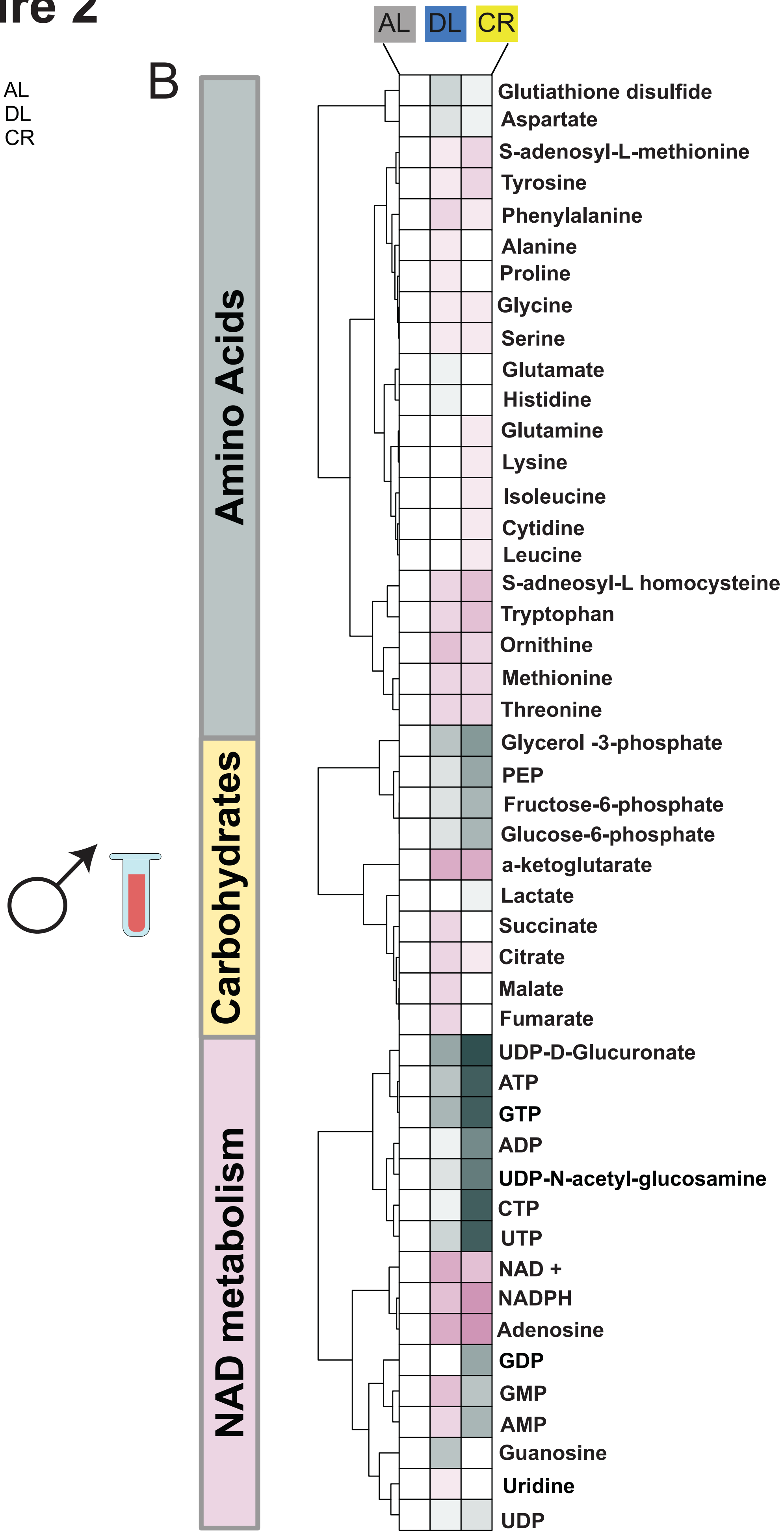

**Supplementary Figure 2: Plasma metabolome profile of male 3xTg mice.**

Targeted metabolomics were performed on the plasma of male 3xTg mice fed AL, Diluted AL and CR diets. (A) Principal Component Analysis (PCA) of plasma metabolites from 3xTg males fed on AL, DL or CR diets (B) Heatmap of 47 targeted metabolites, represented as log2-fold change vs. AL-fed mice. (A-B) n=4 AL, n=3 DL and n=5 CR fed 3xTg biologically independent mice.

### Supplementary Fig 3

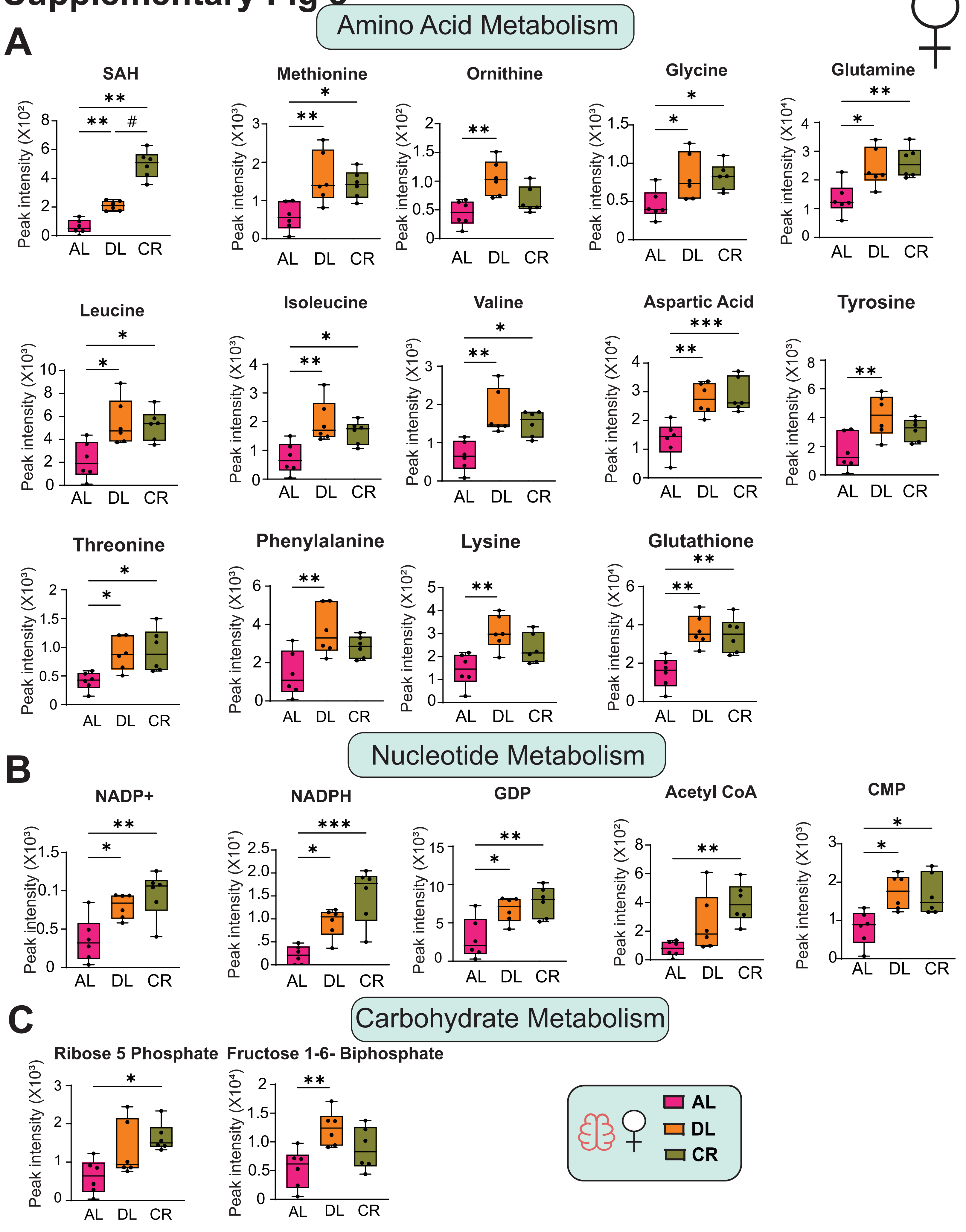

**Supplementary Figure 3: Significantly abundant brain metabolites in female 3xTg-AD mice.**

Targeted metabolomics were performed on the brain of female 3xTg mice fed AL, Diluted AL and CR diets (n = 6) biologically independent mice per diet. (A) Relative abundance of significant amino acid metabolites (B) nucleotide metabolites and (C) carbohydrate metabolites. (A-C) * symbol represents a significant difference versus AL mice ($p \leq 0.05$); # symbol represents a significant difference versus Diluted AL mice ($p \leq 0.05$) based on Tukey's test post one-way ANOVA. Overlaid box plots show center as median and 25th-75thpercentiles; whiskers represent minima and maxima.

### Supplementary Figure 4

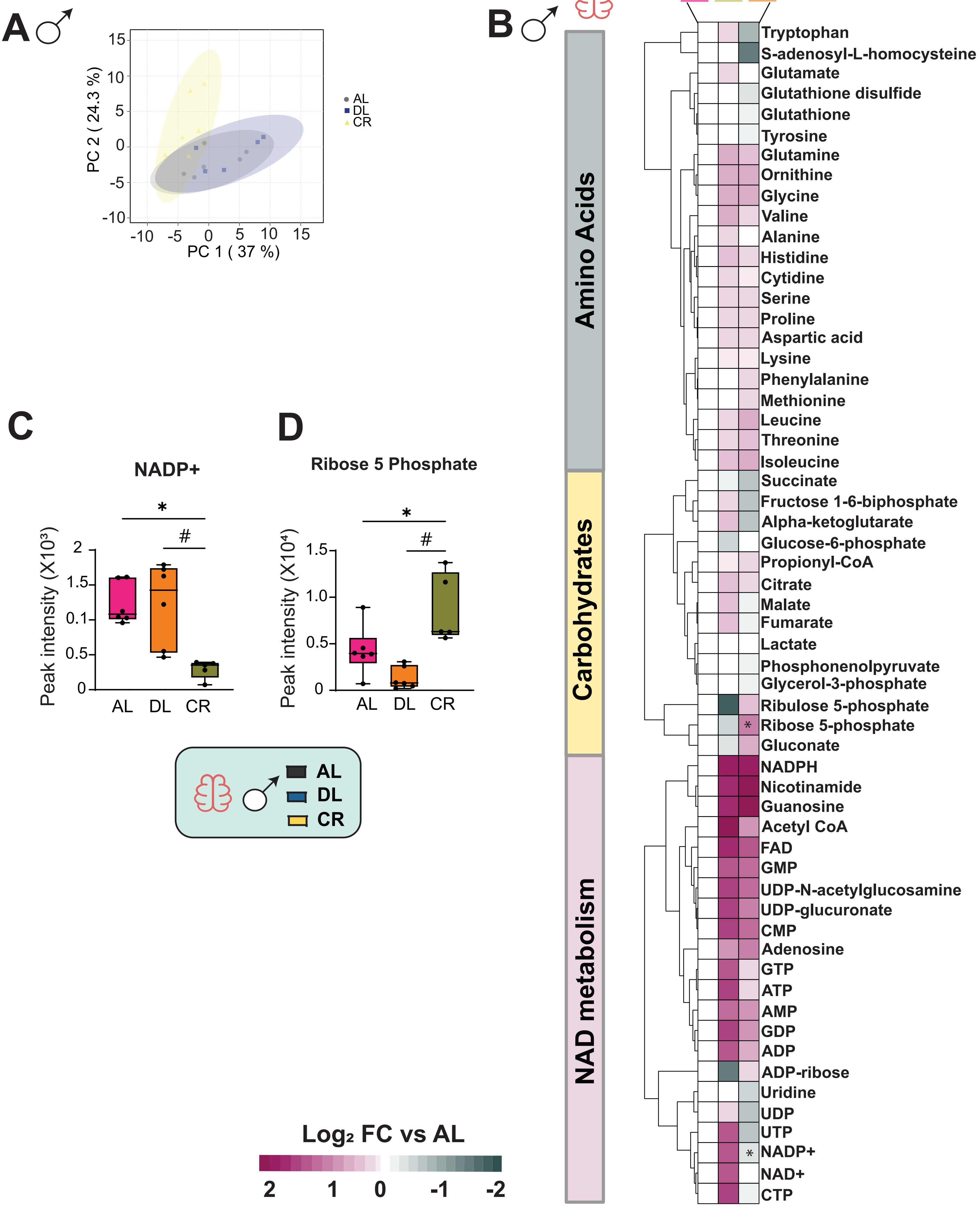

**Supplementary Figure 4: Brain metabolome profile of 3xTg male mice.**

Targeted metabolomics were performed on the brain of male 3xTg mice fed AL, Diluted AL and CR diets. (A) Principal Component Analysis (PCA) of plasma metabolites from 3xTg males fed on AL, DL or CR diets (B) Heatmap of 47 targeted metabolites, represented as log2-fold change vs. AL-fed mice. (C) Relative abundance of nucleotide metabolite NADP+ (D) relative abundance of carbohydrate metabolite Ribose 5 phosphate. (A-D) n=6 AL, n=6 DL and n=5 CR fed 3xTg biologically independent mice. (B-D) * symbol represents a significant difference versus AL mice ($p \leq 0.05$); # symbol represents a significant difference versus Diluted AL mice ($p \leq 0.05$) based on Tukey's test post one-way ANOVA. Overlaid box plots show center as median and 25th-75thpercentiles; whiskers represent minima and maxima.

### Supplementary Figure 5

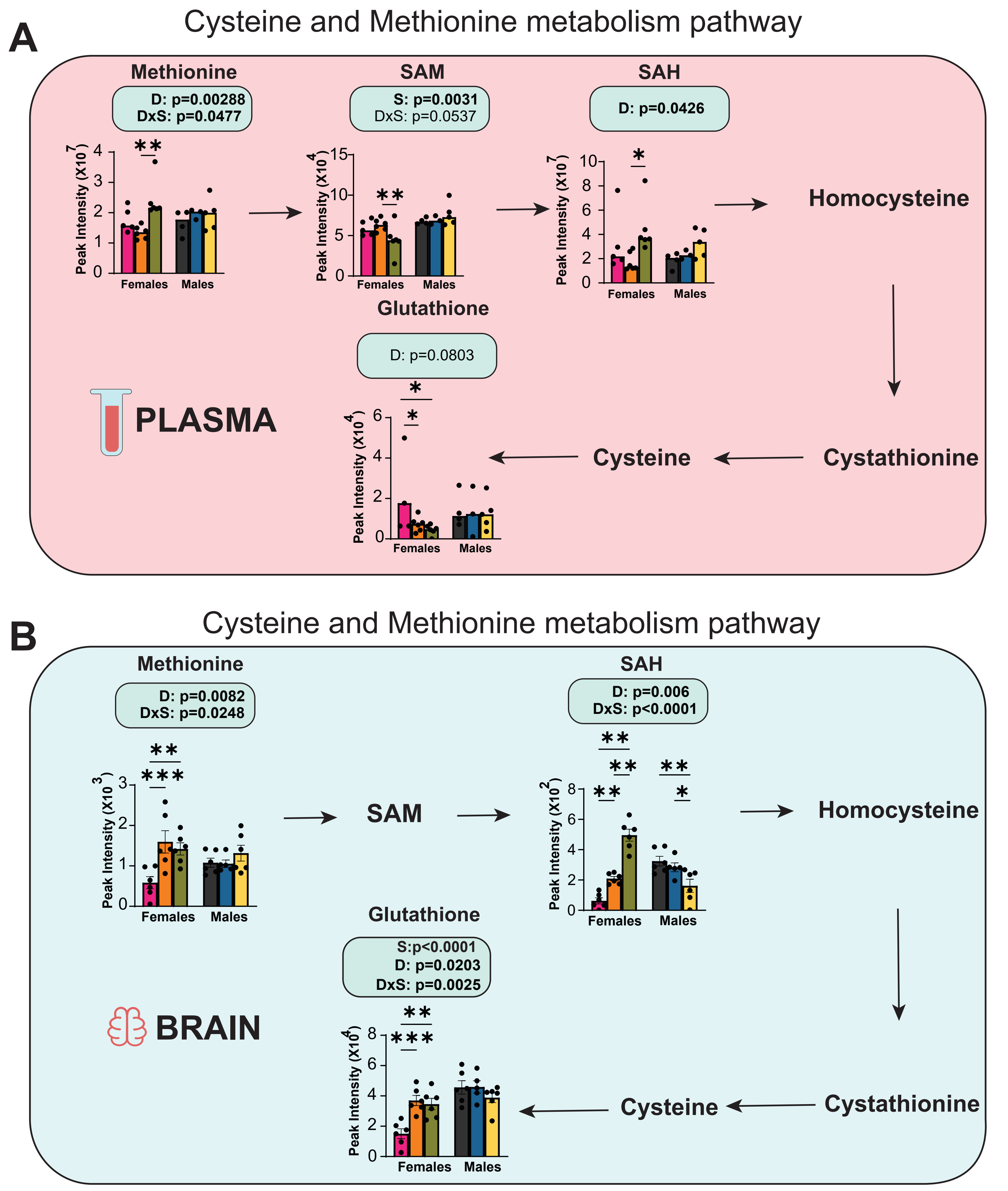

**Supplementary Figure 5. Plasma and brain metabolite data involved in cysteine and methionine metabolism from females and males, related to Figure 4 and Figure 5.**

(A-B) Targeted metabolomics was performed on plasma and brain of both females and males fed on AL, Diluted AL or CR diets. (A) Cysteine and methionine metabolite abundance in plasma from both females and males. (B) Cysteine and methionine metabolite abundance in brain from both females and males. (A-B) For females plasma and brain n=6 biologically independent mice/ group and for males plasma n=4 AL, n=3 DL and n=5 CR and for brain n=6 AL, n=6 DL and n=5 CR fed 3xTg biologically independent mice were used. (A-B) Statistics for the overall effects of diet, sex and the interaction represent the p value from a 2-way ANOVA, *$p<0.05$, from a Sidak's post-test examining the effect of parameters identified as significant in the 2-way ANOVA. Data represented as mean ± SEM.

### Supplementary Figure 6

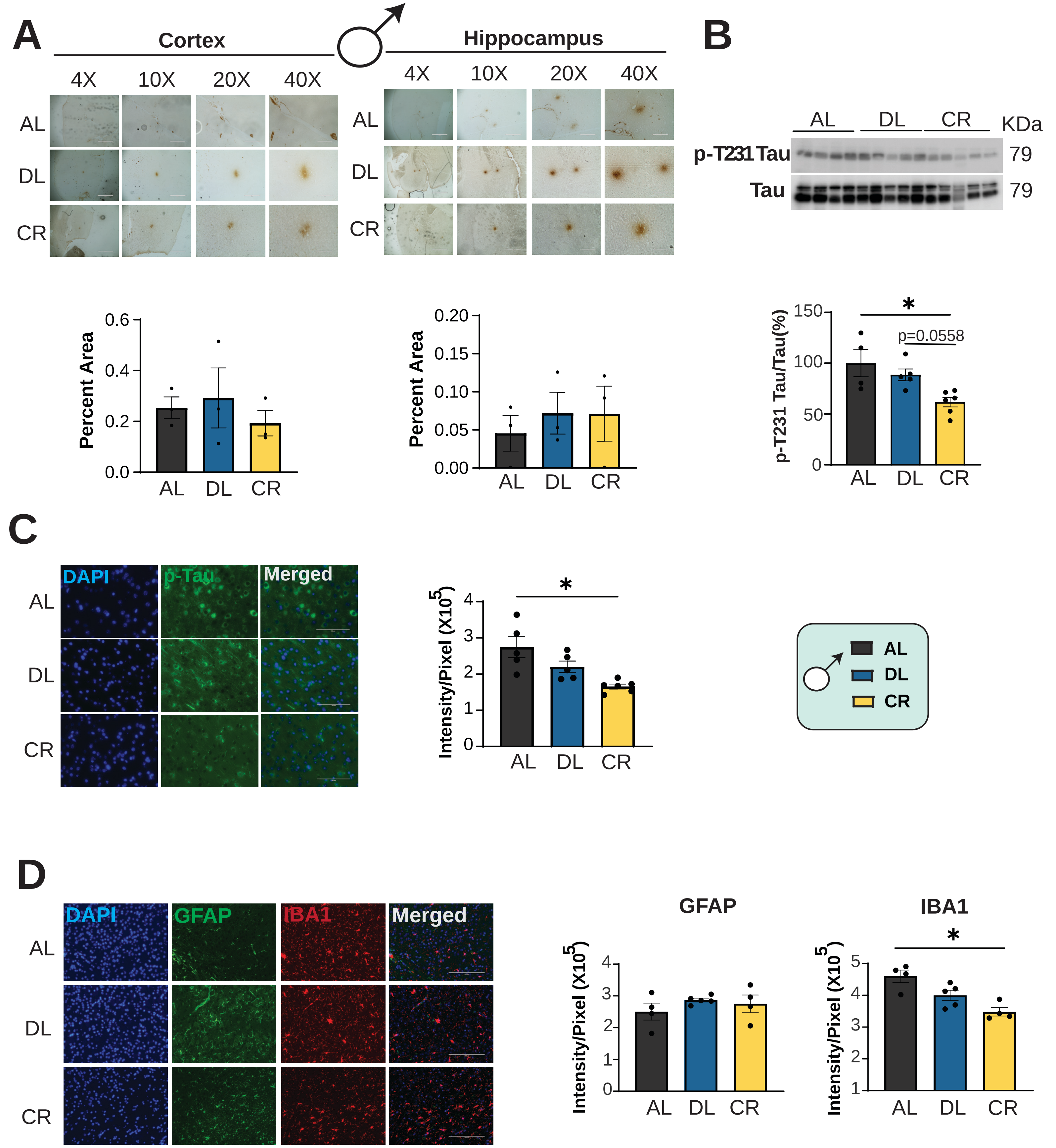

**Supplementary Figure 6: Reduced hyperphosphorylation of tau requires intermeal fasting periods in male 3xTg-AD mice.**

(A-C) Analysis of AD neuropathology in male 3xTg mice fed on AL, Diluted AL or CR diet from 6-15 months of age. (A) Representative plaque images of DAB staining with 6E10 antibody in the cortex and hippocampus of male 3xTg mice. 4x, 10x, 20x and 40x magnification shown; scale bar in the 4x image is 1000 μM, 10x image is 400μM, 20x is 200 μM and 40x is 100 μM. Quantification of plaque area in males shown under each representative blots from cortex and hippocampus respectively. For cortex and hippocampus n=3 biologically independent mice per group. (B) Western blot analysis of phosphorylated Th231 Tau in male 3xTg mice (n=4 AL, n=5 DL and n=5 CR fed 3xTg biologically independent mice. (C) Representative immunofluorescence images in the cortex of 3xTg females stained with p-Tau Thr231 antibody (AT180), 40x magnification shown; scale bar 100 μM. Quantitative analysis of fluorescence intensity. n=5 3xTg biologically independent mice/ group. (D) Immunostaining and quantification of 5 μm paraffin-embedded brain slices for astrocytes (GFAP) and microglia (Iba1) in male 3xTg mice. 20x magnification shown; Scale bar is 200 μM. (C) n=4 AL, n= 5 DL and n=4 CR fed 3xTg biologically independent mice. (A-C) *p<0.05, Tukey post-test examining the effect of parameters identified as significant in the one-way ANOVA. Data represented as mean ± SEM.

**Supplementary Tables**

**Supplementary Table 1:** Diet composition and calorie content for diets used in this study.

**Supplementary Table 2:** List of all 47 metabolites and their compound names identified through targeted metabolomics in the plasma of female 3xTg mice.

**Supplementary Table 3:** $Log_2$ fold-changes from plasma metabolomic analysis in 3xTg female mice shown in **Figure 4B.**

**Supplementary Table 4:** List of all 47 metabolites and their compound names identified through targeted metabolomics in the plasma of male 3xTg mice.

**Supplementary Table 5:** $Log_2$ fold-changes from plasma metabolomic analysis in 3xTg male mice shown in **Supplementary Figure 2B.**

**Supplementary Table 6:** Pathway enrichment analysis of the plasma metabolites in 3xTg mice shown in **Figure 4C**. Significantly up and down regulated pathways for each sex and diet were determined using metabolite set enrichment analysis (MSEA) unadjusted P value $< 0.1$.

**Supplementary Table 7:** List of all 58 metabolites and their compound names identified through targeted metabolomics in the brains of female 3xTg mice.

**Supplementary Table 8:** $Log_2$ fold-changes from brain metabolomic analysis in 3xTg female mice shown in **Figure 5B.**

**Supplementary Table 9:** Pathway enrichment analysis of the brain metabolites in 3xTg mice shown in **Figure 5C**. Significantly up and down regulated pathways for each sex and diet were determined using metabolite set enrichment analysis (MSEA) unadjusted P value $< 0.1$.

**Supplementary Table 10:** List of all 58 metabolites and their compound names identified through targeted metabolomics in the brains of male 3xTg mice.

**Supplementary Table 11:** $Log_2$ fold-changes from brain metabolomic analysis in 3xTg male mice shown in **Supplementary Figure 4B.**

**Supplementary Table 12:** Antibodies used for both western blotting and immunohistochemistry
